# Modular Fluorescent Nanoparticle DNA Probes for Detection of Peptides and Proteins

**DOI:** 10.1101/2021.07.30.454524

**Authors:** Cassandra M. Stawicki, Torri E. Rinker, Markus Burns, Sonal S. Tonapi, Rachel P. Galimidi, Deepthi Anumala, Julia K. Robinson, Joshua S. Klein, Parag Mallick

**Author notes:** These authors contributed equally to this work (Cassandra M. Stawicki, Torri E. Rinker). To whom correspondence should be addressed Corresponding author at: 201 Industrial Rd #310, San Carlos, CA 94070, USA. Email Address (Torri Rinker).

## Abstract

Fluorescently labeled antibody and aptamer probes are used in biological studies to characterize binding interactions, measure concentrations of analytes, and sort cells. Fluorescent nanoparticle labels offer an excellent alternative to standard fluorescent labeling strategies due to their enhanced brightness, stability and multivalency; however, challenges in functionalization and characterization have impeded their use. This work introduces a straightforward approach for preparation of fluorescent nanoparticle probes using commercially available reagents and common laboratory equipment. Fluorescent polystyrene nanoparticles, Thermo Fisher Scientific FluoSpheres™, were used in proof-of-principle studies. Particle passivation was achieved by covalent attachment of amine-PEG-azide to carboxylated particles, neutralizing the surface charge from -43 to -15 mV. A conjugation-annealing handle and DNA aptamer probe was attached to the azide-PEG nanoparticle surface either through reaction of pre-annealed handle and probe or through a stepwise reaction of the nanoparticles with the handle followed by aptamer annealing. Nanoparticles functionalized with DNA aptamers targeting histidine tags and VEGF protein had high affinity (EC_50s_ ranging from 3-12 nM) and specificity, and were more stable than conventional labels. This protocol for preparation of nanoparticle probes relies solely on commercially available reagents and common equipment, breaking down the barriers to use nanoparticles in biological experiments.

## Introduction

Fluorescently labeled antibody and aptamer probes are used to visualize and quantify biological molecules in the fields of biology, chemistry, and biomedicine^1^. They are employed in applications to study receptor-ligand binding in plate-based assays^2^, single-molecule fluorescent microscopy analyses^3^, and flow cytometry^4^ and in applications requiring cell-type specific targeting such as theranostics^5–7^ and *in vivo* imaging^8, 9^. Several characteristics are universally important for fluorescent probes: stability, affinity, brightness, and biocompatibility.

Conventional fluorescent labels such as luminescent metal complexes, proteins, and organic dyes can suffer from insufficient brightness, poor stability, and photobleaching^10–12^. In addition, traditional labels often chemically interact with biomolecules to the detriment of both the fluorescent probe and the biological system^13^.

Nanoparticles offer numerous advantages over conventional labels. First, fluorescent nanoparticles can be much brighter than conventional labels due to their high molar extinction coefficients and quantum yields^14, 15^. Second, nanoparticles have high photostability due to protective encapsulation of fluorescent dyes or to the mechanism of their fluorescence^12, 16^.

Finally, nanoparticles can be coated with biologically inert chemicals like poly(ethylene-glycol) (PEG) to improve biocompatibility^17^. Several types of nanoparticles have been used as fluorescent labels^16, 18, 19^, each with its own unique set of advantages and disadvantages. Dye-doped polymer particles are composed of polymer matrices such as polystyrene or polyacrylate and encapsulate the fluorescent dyes. Polymeric nanoparticles are inexpensive, bright, easy to functionalize, and biocompatible; however, they interact non-specifically if insufficiently passivated and are subject to photobleaching, albeit on far greater time scales than conventional labels^16^. Quantum dots, which are composed of semiconducting materials and are intrinsically fluorescent, have high quantum yields and molar extinction coefficients, are very photostable, and can be modified with exterior organic capping to enhance biocompatibility^20^; however, quantum dots are cytotoxic, limiting their use *in vitro* and *in vivo*^21^. Furthermore, quantum bots blink, which can be a hinderance in single-molecule studies^12^. Evaluation of other nanoparticle types, such as gold, up-converting, carbon, and silica nanoparticles, as fluorescent labels is ongoing^18^.

Although fluorescent nanoparticles may offer improvements to conventional labels, challenges in synthesis, functionalization, and passivation have limited their use. For example, incorporating dyes into silica nanoparticles or polymeric matrices requires a detailed understanding of fluorophore quenching behavior and expertise in emulsion and polymerization techniques^16, 19, 22^. The synthesis of quantum dots requires specialized equipment for techniques like e-beam lithography, formation of microemulsions, and sputtering for vapor-phase synthesis^23, 24^. Once synthesized, nanoparticles require specialized equipment for characterization, such as electron microscopy, not available to every laboratory^16, 25^. After synthesis, nanoparticles are functionalized by attaching probes to the surface, an often arduous and expensive process that requires specific expertise^26, 27^. For example, in the case of silica nanoparticles, a reaction with (3-aminopropyl)triethoxysilane (APTES) is often used to introduce a biocompatible amine-functionality to the particle, but this reaction must be controlled to avoid APTES polymerization and can require additional chemical modification to allow compatibility with probe functional groups^28, 29^. In addition to functionalizing the nanoparticle with an active chemistry, the affinity reagent probe must be functionalized with a compatible conjugation- annealing handle. Reagents linked to common chemical moieties like thiols and amines are readily available from commercial vendors, but specialized and often more desirable functional groups require custom synthesis and characterization. Finally, achieving sufficient passivation on the nanoparticle surface can be technically challenging. For instance, quantum dots must be capped with hydrophilic materials, like amphiphilic polymers, to improve solubility and colloidal stability in aqueous solutions^23, 30^. For polymeric particles, passivation prevents undesirable interactions with the biological environment. Moreover, the passivated layer provides functional groups for further attachment of probes^16^. Although individual solutions to nanoparticle challenges have been identified, no common set of best practices are available.

To simplify the preparation and use of nanoparticle-based probes, we set out to develop a protocol for functionalizing off-the-shelf reagents to generate fluorescent nanoparticle probes. For proof-of-principle experiments, commercially available dye-laden polystyrene nanoparticles (FluoSpheres™, ThermoFisher) and quantum dots (Qdot™ 655 ITK™ Carboxyl Quantum Dots, ThermoFisher) were chosen due to their stability, brightness, and biocompatibility. These nanoparticles were modified with PEG to improve affinity and colloidal stability. Finally, aptamer probes were attached to the particle by annealing disparate probes to a common oligonucleotide conjugated to a chemical moiety for covalent attachment to the nanoparticle; we call this oligonucleotide the conjugation-annealing handle. This modular design allows nanoparticle to be easily functionalized with a new probe. In our proof-of-concept studies, we demonstrated the utility of our method by functionalizing FluoSphere and Qdot fluorescent nanoparticles with previously described DNA aptamer probes^31, 32^. Our studies show that our fluorescent nanoparticle probes meet the stability, affinity, brightness, biocompatibility, modularity, and reagent availability requirements necessary for a fluorescent label. We provide a supplemental protocol with easy-to-follow instructions so that other researchers can create their own fluorescent nanoparticle probes.

## Results

### Fluorescent nanoparticle probe fabrication

Our method consists of three steps: passivation with chemically active PEG, attachment of a conjugation-annealing handle to enhance modularity, and attachment of a DNA-based aptamer probe (**Figure 1**). All materials used in this fabrication method are commercially available (**Table 1**). For initial protocol optimization and characterization, 40-nm and 200-nm yellow-green or dark-red FluoSpheres, which are nanoparticles encapsulating fluorescent dyes within a polymeric polystyrene matrix, were used as the nanoparticle label. To attach the passivating PEG-layer, the FluoSpheres were activated with 1-ethyl-3-(3- dimethylaminopropyl)carbodiimide/ *N*-hydroxysuccinimide (EDC/NHS) and reacted with PEG functionalized with an amine and an azide moiety for DNA conjugation. We attached the conjugation-annealing handle and DNA aptamer probe to the nanoparticle either through a single reaction between the nanoparticle and the pre-annealed probe complex or through a stepwise reaction with conjugation of the handle to the nanoparticle followed by annealing of the aptamer (**Figure 1A**). The handle consists of a 3’ dibenzocyclooctyne (DBCO) for conjugation to the PEG layer and either a 24-base-pair annealing sequence (Illumina P7 primer sequence^33^) or a poly-T sequence (21 thymidine bases; **Figure 1B**). The aptamer includes a complementary sequence that anneals to the handle. The optimized protocol is included in the Supplementary Information.

**Figure 1.**
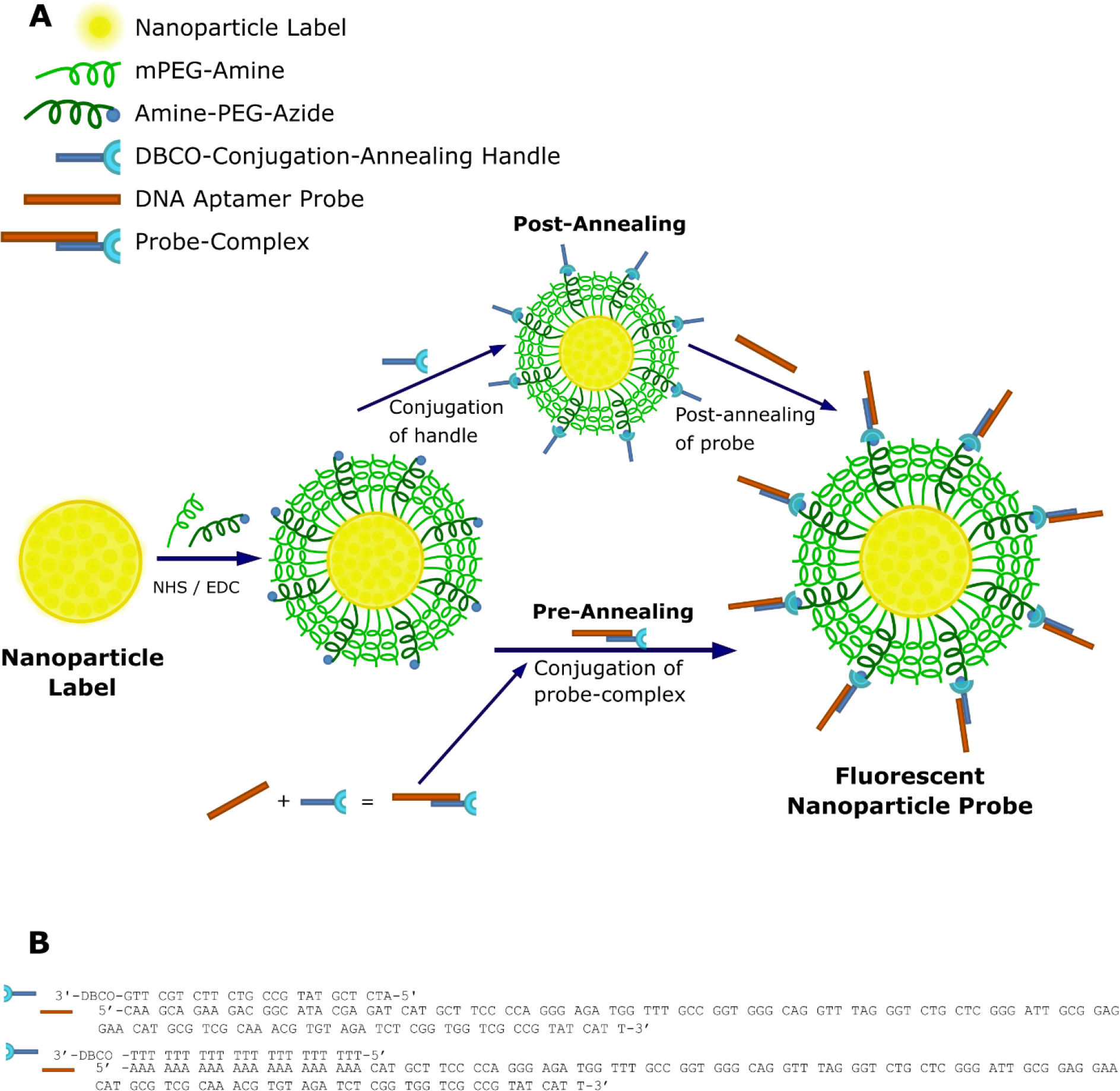
Fluorescent nanoparticle probe fabrication and DNA aptamer probe attachment. (A) Carboxylate-modified nanoparticle labels are PEGylated for passivation and click chemistry modification using a combination of mPEG-amine and amine-PEG-azide and NHS/EDC chemistry. DNA aptamer probes are attached to the particles via annealing to the conjugation-annealing handle either before or after it is conjugated to the PEG layer. Figure is not to scale. (B) The conjugation-annealing handle is used to conjugate a DNA aptamer probe to the nanoparticle. The same handle can be used to conjugate different aptamer probes. The handle consists of a DNA sequence complementary to a short region on the 5’ end of the DNA aptamer probe and a DBCO functional group on the 3’ end for covalent attachment to the particle. The DNA aptamer probe consists of a variable probe region that can bind to a target of interest and an annealing region that allows attachment to the conjugation-annealing handle.

**Table 1.** Commercially Available Reagents for Fluorescent Nanoparticle Probes

| Component of Fluorescent Nanoparticle Probe | Vendor | Catalog No. | Product Name |
| --- | --- | --- | --- |
| Nanoparticle label | Thermo Fisher: Molecular Probes | F8795 | FluoSpheres™ Carboxylate-Modified Microspheres, 0.04 μm, yellow-green fluorescent (505/515), 5% solids, azide free |
|  | Thermo Fisher: Molecular Probes | F8811 | FluoSpheres™ Carboxylate-Modified Microspheres, 0.2 μm, yellow-green fluorescent (505/515), 5% solids |
|  | Thermo Fisher: Molecular Probes | F8807 | FluoSpheres™ Carboxylate-Modified Microspheres, 0.2 μm, dark-red fluorescent (660/680), 5% solids |
|  | Thermo Fisher: Molecular Probes | Q21321MP | Qdot™ 655 ITK™ Carboxyl Quantum Dots |
| Passivation and Functionalization | Creative PEGWorks | PHB-1882 | Amine-PEG-azide, MW 2 kDa |
|  | Creative PEGWorks | PLS-269 | mPEG-Amine, MW 2 kDa |
| Conjugation Annealing Handle | Integrated DNA Technologies | Custom sequence; Table 3 | <custom_sequence>/iSp9//3DBCON/, HPLC purification |
| DNA Aptamer Probe | Integrated DNA Technologies | Custom sequence; Table 3 | 20 nmole Ultramer |

### Optimization of nanoparticle PEGylation

To demonstrate that methoxy-PEG (mPEG) was effectively conjugated to the nanoparticles, we first introduced either 1.6-kDa mPEG-amine (reactive), methoxy-PEG-OH (unreactive), or buffer only at a concentration of 10^8^ PEG molecules/particle and examined the impact of PEG conjugation on nanoparticle zeta potential. Zeta potential measurements rely on scattering of incident laser light to measure the velocity of particle motion under application of an electric field^34^. Changes in zeta potential of reacted particles indicate consumption of highly negatively charged carboxylate groups during conjugation and charge-shielding of any remaining negative charge by the PEG layer^35^, which should only occur if chemical conjugation has occurred. Zeta potentials of -43 mV in the buffer-treated and mPEG-OH-treated samples indicated minimal non-specific interactions of PEG with the activated nanoparticle. The zeta potential was -15.4 mV when nanoparticles were incubated with 10^8^ mPEG-amine molecules/particle (**Figure 2A**), indicating that PEGylation occurred via covalent linkage to carboxylate-modified nanoparticles.

**Figure 2.**
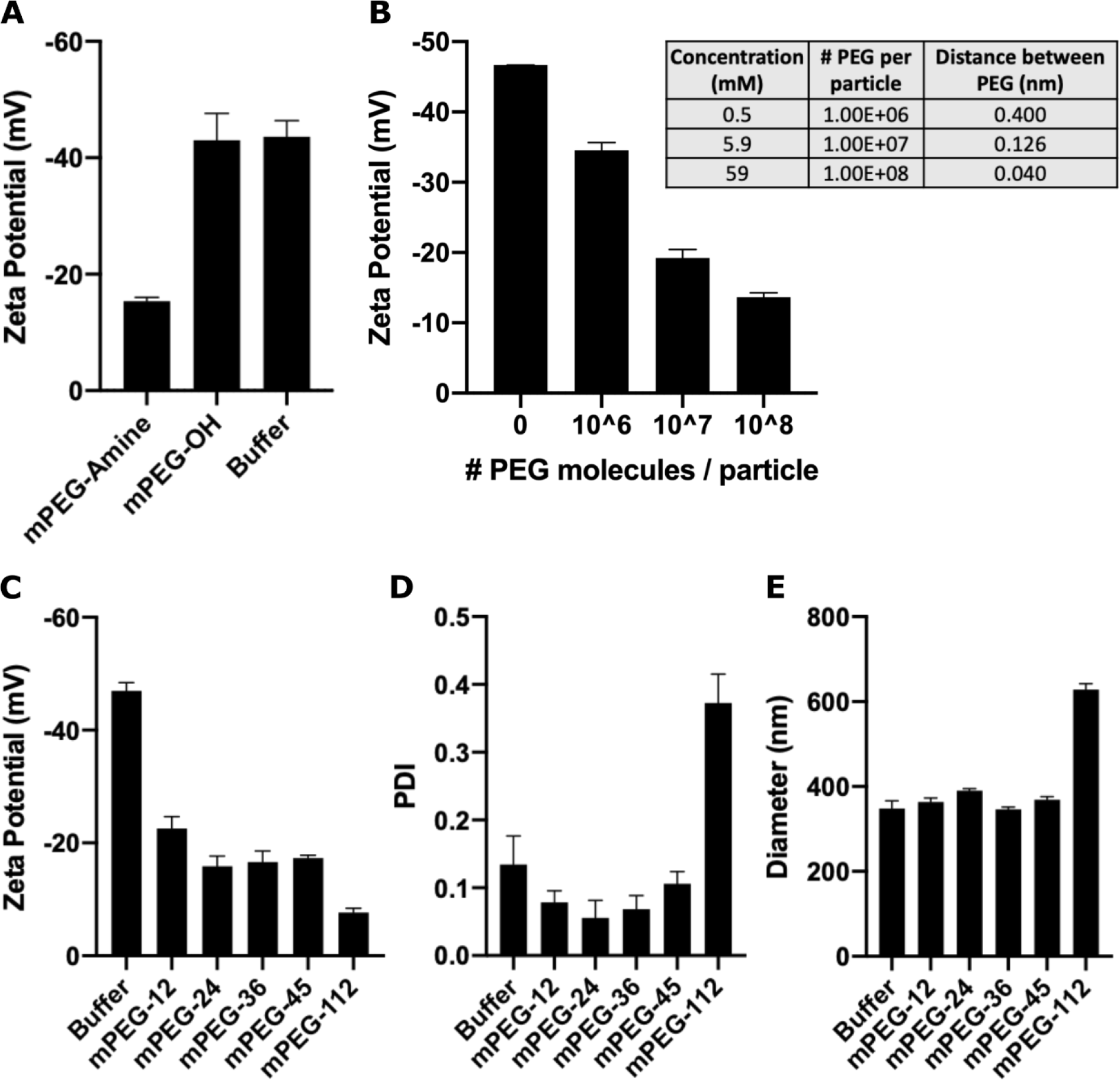
PEGylation of carboxylate-modified FluoSpheres™. (A) Zeta potential measurements for carboxylated FluoSpheres™ activated with NHS/EDC and reacted with mPEG-amine, mPEG-OH, or buffer only. Data are means ± standard deviation of three measurements per sample for one of two independent experiments, analyzed in duplicate. (B) Zeta potential measurements for carboxylated FluoSpheres™ activated with NHS/EDC and reacted with increasing concentrations of mPEG-amine. Data are means ± standard deviation of one experiment, analyzed in triplicate. Table lists theoretical distances between PEG molecules at each concentration. (C-E) FluoSpheres™ were conjugated with PEGs of indicated molecular weights, and C) zeta potentials, D) PDIs, and E) hydrodynamic diameters were determined. Shown are means ± standard deviation of three measurements per sample, n=2-3.

Next, we tested a range of PEG grafting densities by evaluating the zeta potential of nanoparticles reacted with 10^6^ to 10^8^ PEG molecules/particle. Increased PEG density reduced the magnitude of negative nanoparticle surface charge from -35 mV to -14 mV, suggesting a more highly passivated particle at higher PEG densities (**Figure 2B**).

We also evaluated the impact of PEG molecular weight on nanoparticle surface charge and colloidal stability. Colloidal stability was inferred by dynamic light scattering (DLS). In DLS, particle velocity is measured by subjecting the nanoparticle suspension to an incident laser and measuring the constructive and destructive interference patterns produced by scattered light over time, which can then be used to calculate the hydrodynamic radius of the nanoparticles^34, 36^. In general, PEGs with molecular weights of about 2 kDa (PEG-45) or larger are used for nanoparticle passivation^37^, so we tested PEGs ranging from PEG-12 to PEG-112. As the molecular weight of PEG increased, nanoparticle surface charge neutralization increased, indicative of more complete passivation. PEG-112, the longest polymer tested, had the greatest decrease from an activated zeta potential of -46 mV to a final zeta potential of -7.75 mV (**Figure 2C**). PEG-112-conjugated particles had a polydispersity index (PDI) of 0.36 and hydrodynamic diameter of 628 nm; particles conjugated with lower molecular weight PEGs had PDIs of 0.04-0.1 and diameters of 332-394 nm (**Figure 2D and E**). As PDIs greater than 0.1 suggest nanoparticle aggregation^34^, PEG-112 was eliminated from further study. Particles conjugated to PEG-45 had a zeta potential of -17 mV, a PDI of 0.11 and diameter of 369 nm, indicating that passivation was achieved and colloidal stability was maintained. Thus, PEG-45 was used for all subsequent studies at a concentration of 3.5x10^7^ PEG molecules per 200-nm particle and 1.4x10^5^ PEG molecules per particle for 40-nm nanoparticles (**Figure S1**).

We hypothesized that a higher percentage of azide-PEG-amine would allow conjugation of more probes per particle, which would result in higher affinity of the nanoparticle probe to its target. We compared mPEG-amine:azide-PEG-amine ratios of 75:25, 95:5, 99.5:0.5, 99.95:0.05, and 99.995:0.005 in protein binding studies. Similar on-target binding was observed at the 75:25 and 95:5 ratios, while lower binding was observed for ratios with less azide-PEG-amine (**Figure S2**). To achieve the highest binding possible while conserving reagents, we selected a ratio of 95:5 mPEG-amine:azide-PEG-amine for the remainder of these studies.

### Quantification of DNA aptamer probe-to-nanoparticle ratio

Next, we evaluated changes in particle charge and size with addition of the conjugation- annealing handle and of the DNA aptamer probe. The addition of the handle to the nanoparticle surface increased the overall negative charge on the nanoparticle from -13 mV to -19 mV, and subsequent annealing of the DNA aptamer probe further increased negative charge to -21 mV (**Figure 3A**). Nanoparticle diameter increased as each component was added from 335 nm for the PEGylated particle to 367 nm for fully functionalized fluorescent nanoparticle probes (**Table 2**). The PDI remained close to 0.1, indicating that colloidal stability was maintained. The decrease in surface charge and slight increase in size indicated successful attachment of both the conjugation-annealing handle and the probe to the nanoparticle.

**Figure 3.**
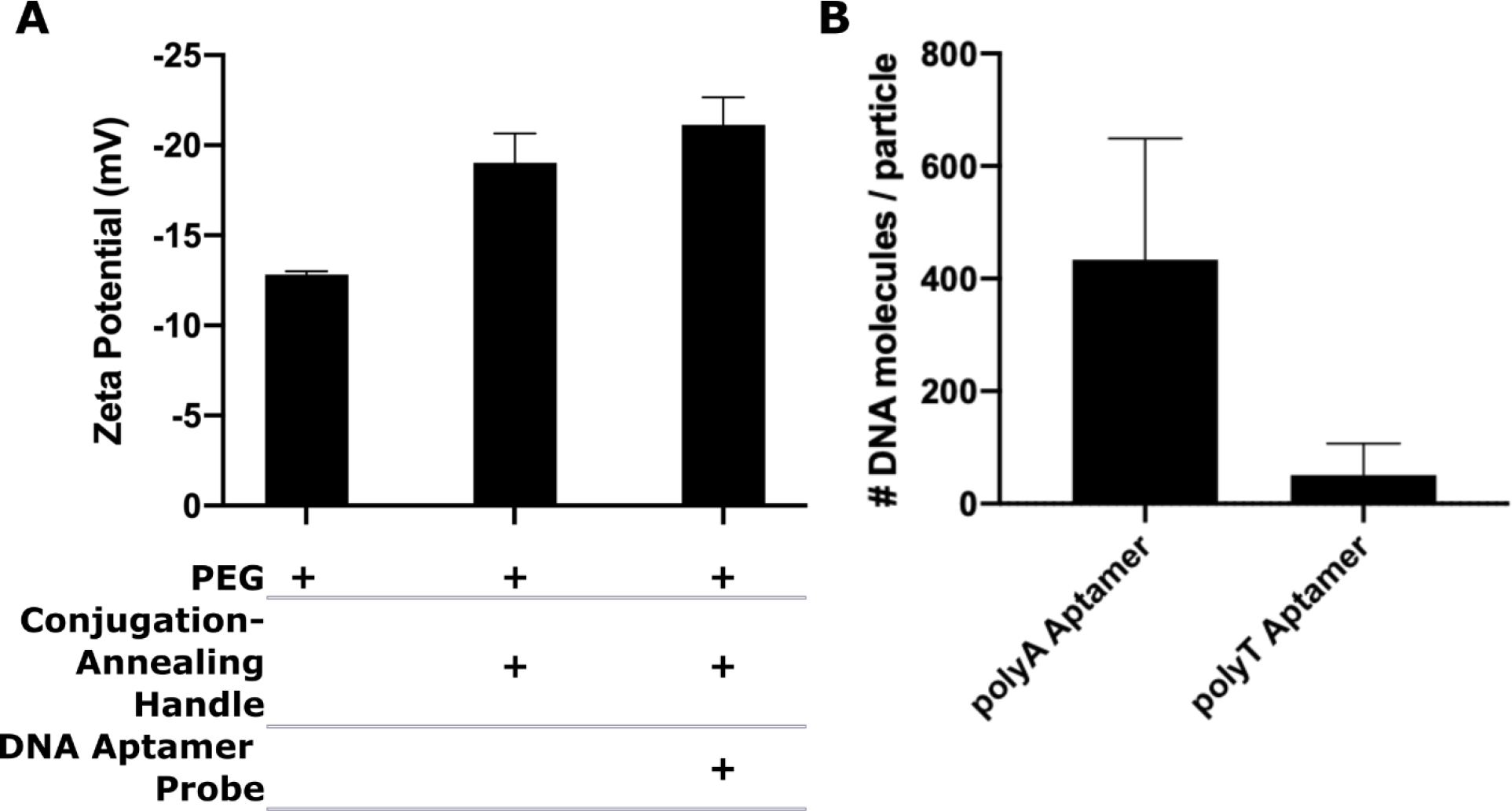
Characterization of DNA aptamer probe attachment to PEGylated nanoparticle. (A) Zeta potential measurements after PEGylation, conjugation of conjugation-annealing handle, and annealing of DNA aptamer probe. Data are means ± standard deviation of one experiment, analyzed in triplicate. (B) qPCR quantification of aptamer numbers for particles functionalized with complementary polyA aptamer or non-complementary (negative control) polyT aptamer. Data shown are means ± standard deviation, n=3.

**Table 2.** Summary of particle diameter, PDI, and zeta potential of EDC/NHS activated carboxylate-modified FluoSpheres™, PEGylated FluoSpheres™ and fluorescent nanoparticle probes of a 200-nm diameter. Data are shown as means ± standard deviation.

| **Stage in Nanoparticle Fabrication** | **Particle Diameter (nm)** | **Polydispersity index (PDI)** | **Zeta Potential (mV)** |
| --- | --- | --- | --- |
| Activated Nanoparticle | 335 ± 3.0 | 0.13 | -46.9 ± 1.0 |
| PEGylated Nanoparticle | 367 ± 6.0 | 0.12 | -17.3 ± 0.3 |
| Fluorescent Nanoparticle Probe | 367 ± 0.1 | 0.12 | -21.1 ± 1.5 |

Next, we quantified aptamer attachment to the nanoparticles using qPCR, a technique commonly used to measure the quantity of DNA on a nanoparticle^38^. Amplification was performed of oligonucleotides removed from fabricated nanoparticles by heat. The number of DNA molecules was determined by comparing cycle threshold (Ct) values of the experimental group to a standard curve of the DNA aptamer. About 430 DNA aptamer probes were detected per 200 nm particle (**Figure 3B**). For particles treated with a negative-control aptamer, which did not have sequence complementary to the conjugation-annealing handle, about 50 aptamers were detected per particle.

In addition, we quantified the number of conjugation-annealing handles/nanoparticle by reacting 40 nm PEGylated particles with a fluorescently labeled conjugation-annealing handle. The number of oligonucleotides per particle increased with the concentration of input conjugation-annealing handle and no attachment was observed for the non-DBCO containing control (**Figure S3**). In addition, the ratio of 20 conjugation-annealing handles per 40 nm particle aligns well with the ratio of 430 aptamers per 200 nm particles when differences in surface area are accounted for. Overall, these data indicate successful aptamer attachment and negligible non-specific binding between non-specific oligonucleotides and nanoparticles.

### Assessment of fluorescent nanoparticle probe binding

We used a plate-based binding assay to determine whether the addition of the label to the probe altered the binding affinity of the probe and to interrogate the differences between several probe attachment methods. In these proof-of-principle studies, we used the B1 aptamer, which binds histidine tags^31^, as our probe. A biotinylated B1 probe detected with traditional detection methods (amplification of signal by streptavidin HRP or conjugation to a fluorescently labeled streptavidin conjugated) bound histidine-histidine-histidine (HHH) with an observed equilibrium dissociation constant (Kd) of 55 nM (**Figure S4A and B**), comparable to the reported Kd of 120 nM^31^.

We evaluated specific and non-specific binding of the fluorescent nanoparticle probes against on-target (HHH peptide, His-tagged Her2 protein) and off-target (Ecoli Lysate, PNG peptide, and Myoglobin) biomolecules. When we evaluated binding of the nanoparticle probes to his-tagged Her2 protein, we observed a 2-fold higher binding by the B1 probe than by the nanoparticles functionalized with a negative control aptamer and observed no binding to Ecoli Lysate (**Figure 4A**). Similar results were seen for off-target binding to myoglobin (**Figure 4B**). To further investigate non-specific binding, we conducted the assay with HHH and with proline-asparagine-glycine (PNG), a peptide that the B1 aptamer does not recognize. Streptavidin AlexaFluor™ 647 Conjugate, a conventional label, was used as a control to ensure that specific and non-specific binding were not due to the label type. Nanoparticles were functionalized with the B1 (on-target) and Her2^39^ (off-target) aptamers to differentiate non-specific binding by the aptamer versus non-specific binding due to the functionalized nanoparticle. Both the Streptavidin AlexaFluor™ 647 Conjugate and fluorescent nanoparticle B1 probe bound to HHH, but we observed minimal non-specific binding to PNG (**Figure 4C**). Nanoparticles functionalized with the Her2 aptamer did not bind detectably to either HHH or PNG. Taken together, these results provide evidence that both particle passivation and probe attachment were successful.

**Figure 4.**
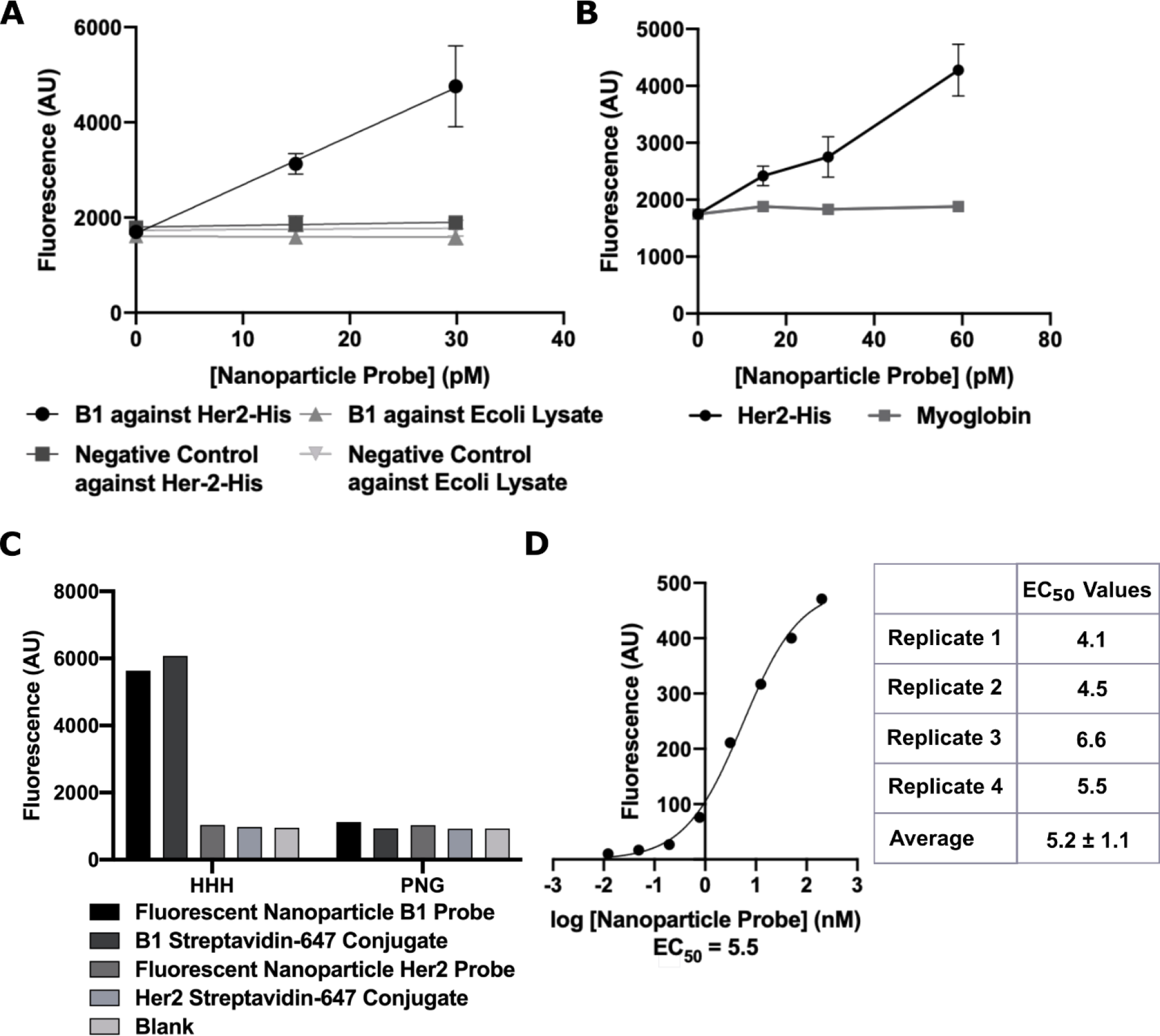
Fluorescent nanoparticle B1 probe binds specifically to HHH peptide and his-tagged protein. (A) Fluorescence signals as a function of concentration of nanoparticle B1 probe and negative control probe upon binding to his-tagged Her2 protein (on-target) and Ecoli lysate (off-target). Data are presented as means ± standard deviation, n=3. (B) Signal from nanoparticle B1 probe against his-tagged Her2 (on-target) and myglobin (off-target). Data are presented as means ± standard deviation, n=3. (C) Fluorescence signals from fluorescent nanoparticle B1 and Her2 probes and B1 and Her2 Streptavidin 647 conjugates against HHH (on-target) or PNG (off- target) peptides. Data is from a single experiment. (D) A representative binding curve of the B1 probe against his-tagged Her2 used to calculate EC50 values. The table shows EC50 values obtained from 4 replicates run across two experiments and reports mean ± standard deviation.

Finally, we evaluated a concentration titration of fluorescent nanoparticle B1 probes against the his-tagged Her2 protein. The half maximal response concentration, EC50, was an average of 5.2 nM (**Figure 4D**). This EC50 value is lower than what was previously reported (120 nM)^31^ and what we found experimentally with other conventional labels (55 nM; **Figure S4B**). We hypothesize that this apparent increase in affinity could result from avidity effects, as described in the Discussion Section.

### Attachment of alternate aptamers to fluorescent nanoparticles

To demonstrate modularity of the fluorescent nanoparticle probe system, nanoparticles were functionalized with H3T, a truncated version of the B1 aptamer^31^, and a VEGF aptamer^32^. These aptamers bind their targets when they are detected with more traditional methods (**Figure S4B and C**). We tested the binding of the HHH peptide by the nanoparticle H3T probe using a plate-based assay. The nanoparticles bound to the peptide target with an average EC50 of 12 nM (**Figure 5A**), stronger affinity than the EC50 of 30 nM found using a more traditional detection approach (**Figure S4B**) and that reported in literature (120 nM)^31^. Nanoparticle probes functionalized with the VEGF aptamer bound VEGF protein and not an off-target protein with an average EC50 of 5.5 nM (**Figure 5B**). This demonstrates the modularity of our probe attachment approach.

**Figure 5.**
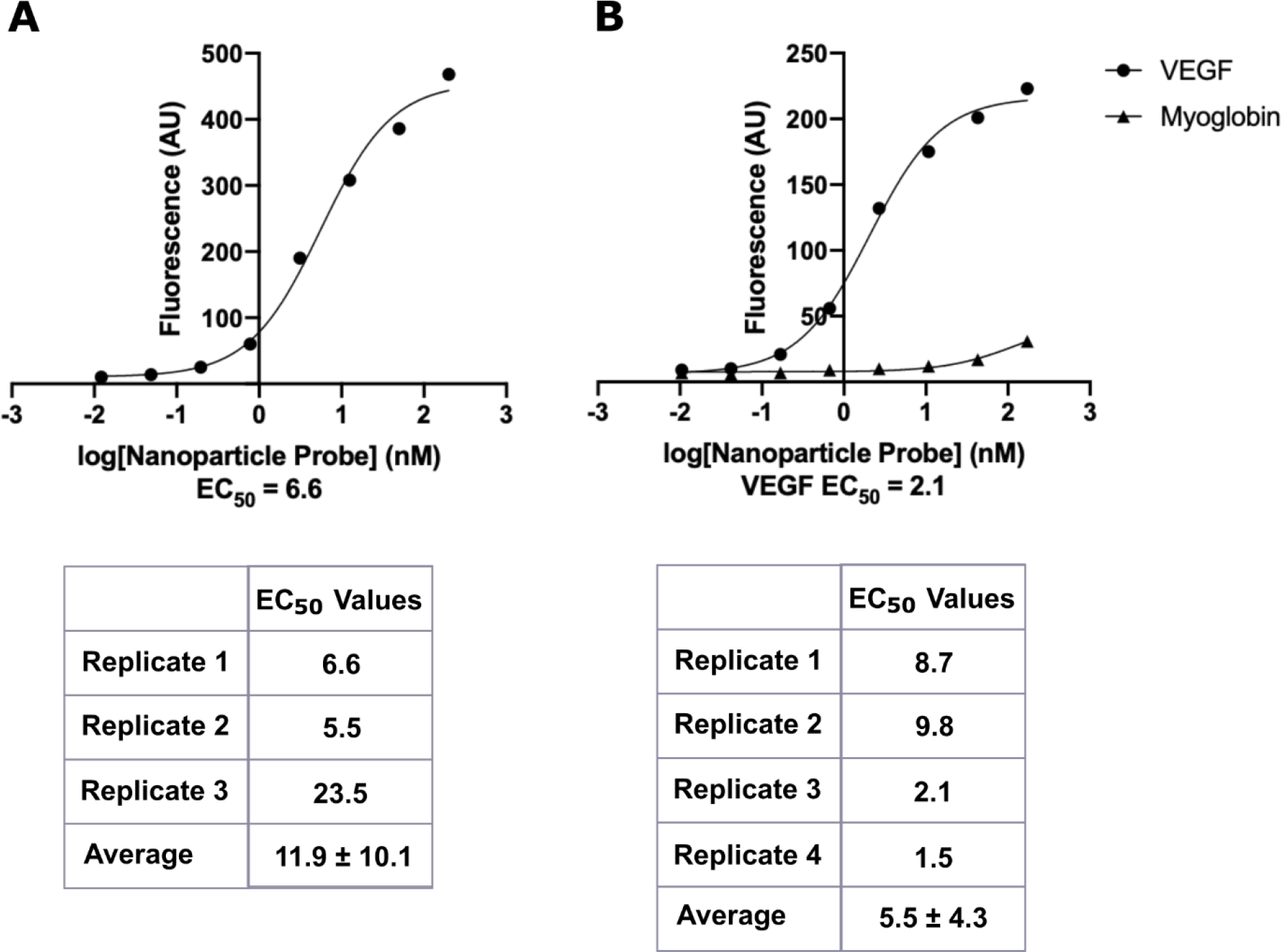
Fluorescent nanoparticles functionalized with various aptamers bind specifically. (A) Representative binding curve for fluorescent nanoparticle H3T probe against HHH peptide. The table shows EC50 values obtained from 3 replicates run across two experiments and reports mean ± standard deviation. (B) Representative binding curve for fluorescent nanoparticle VEGF probe against VEGF (on-target) and myoglobin (off-target). The table shows EC50 values obtained from 4 replicates run across two experiments and reports mean ± standard deviation.

### Alternative attachment strategies

Finally, we evaluated the on-target binding of fluorescent nanoparticle probes fabricated using pre-annealing or post-annealing of the aptamer to the nanoparticle (**Figure 1A**). Pre-annealed and post-annealed fluorescent nanoparticle B1 probes were used to detect an on-target his-tagged protein in a plate-based binding assay. Both pre- and post-annealed particles detected the his-tagged protein with stronger affinity than the off-target myoglobin protein, but at the highest probe concentrations tested, the pre-annealed particles showed 2-fold higher binding to his-tagged protein than did the post-annealed particles (**Figure S5**). To explain these differences, further experimentation is required, but we hypothesize that the concentration of probe mixed with the particles in the post-annealing protocol was not high enough to fully saturate available conjugation-annealing handles.

In addition to annealing the probe to the particle, we evaluated a more traditional attachment approach by directly conjugating a DBCO-modified aptamer to the PEG-azide layer. We evaluated pre-annealed particles and the directly conjugated particles in plate-based binding assays against his-tagged protein. Although both approaches resulted in higher on-target protein binding than off-target binding, the probe prepared using the pre-annealing approach had on-target binding that was about 2-fold greater than the probe prepared using direct conjugation approach (**Figure S6**). Additional studies will be required to determine the reason for this effect. These results indicate that both pre- and post-annealing of aptamers to nanoparticles can be used to achieve specific binding and that annealing approaches are as good as or better than traditional conjugation approaches. The pre-annealing fabrication was used in subsequent experiments.

### Comparison to alternative commercial labeling options

To understand how our nanoparticle labels compare to commercially available labels, we conducted a series of studies to compare affinity, stability, and brightness of the fluorescent nanoparticle probe to off-the-shelf labels including Streptavidin AlexaFluor™ 647 Conjugate, Streptavidin APC Conjugate, and Streptavidin SureLight™ APC. In a plate-based binding assay, we compared the EC50 of our fluorescent nanoparticle B1 probe to commercially available labels attached to biotinylated B1 aptamer (**Figure 6A**). The fluorescent nanoparticle probe had binding affinity approximately equivalent to that of Streptavidin SureLight™ APC, with average EC50s of 2.7 and 9.9 nM, respectively. Streptavidin AlexaFluor™ 647 and Streptavidin APC Conjugates had average EC50 values of 43 nM and 28 nM, respectively. The total number of binding sites per label may impact observed binding affinity due to avidity effects. Information from the vendor indicates that Streptavidin AlexaFluor 647 Conjugate and the Streptavidin APC Conjugate consist of one streptavidin per label, resulting in approximately three biotinylated probe binding sites per label. The Streptavidin SureLight APC consists of multiple streptavidin and APC molecules per conjugate, and conversations with the vendor indicated that the total number of binding sites can vary batch-to-batch. Although more studies will be required to fully understand the observed results, we conclude that the affinity of the fluorescent nanoparticle B1 probe is equal to or greater than commercially available labels.

**Figure 6.**
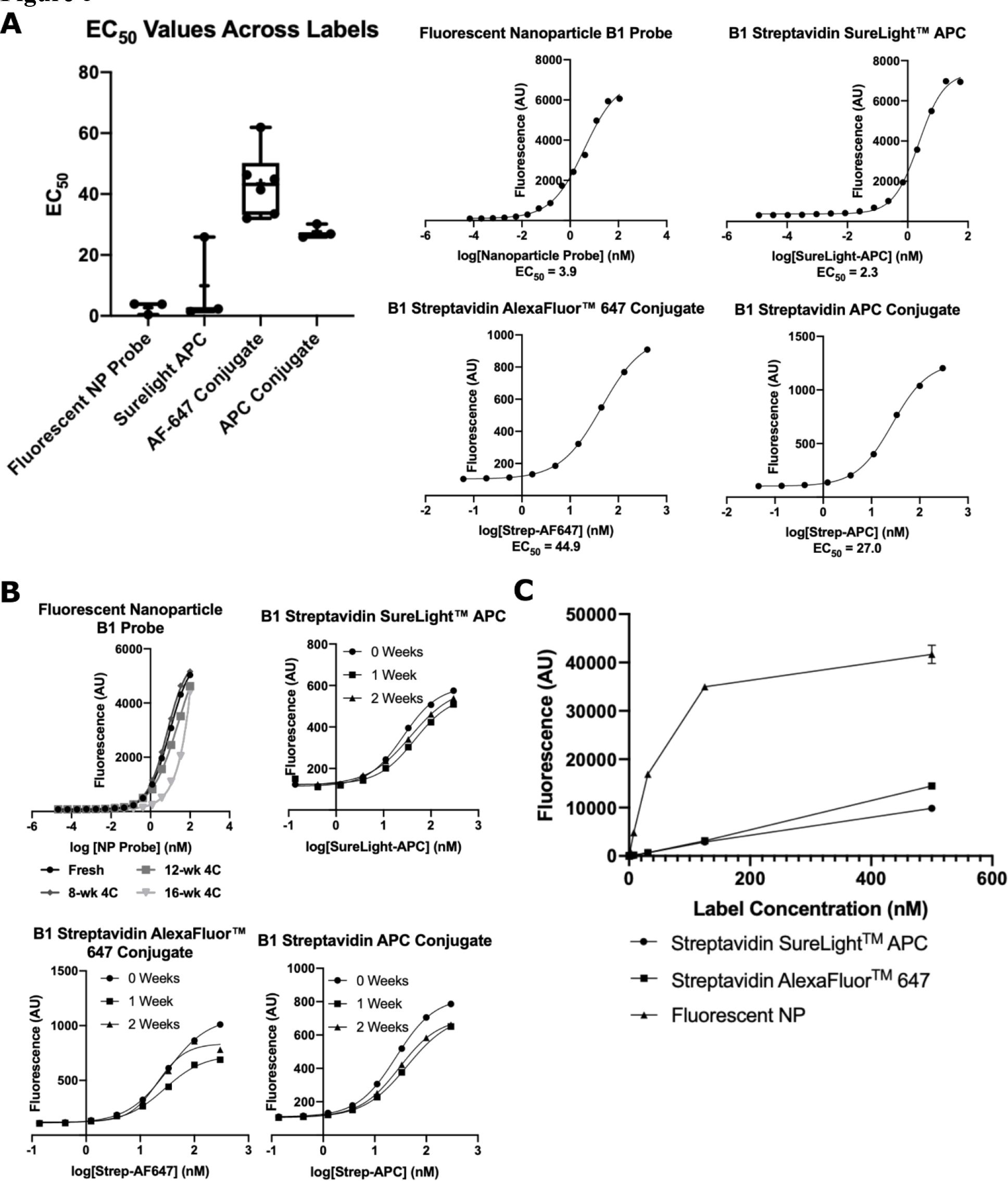
Fluorescent nanoparticle probes compare favorably to commercially available labels. (A) Comparison of EC50 values for fluorescent nanoparticle B1 probe (“Fluorescent NP Probe”), B1 Streptavidin SureLight™ APC (“Surelight APC”), B1 Streptavidin AlexaFluor™ 647 Conjugate (“AF-647 Conjugate”), and B1 Streptavidin APC Conjugate (“APC Conjugate”) against HHH targets. Representative binding curves are shown at the right. Data is shown as box and whisker plots, where the box extends from the 25^th^ to 75^th^ percentiles, the middle line represents the median, the “+” represents the mean, and the whiskers show max and min values. Each point represents an EC50 from an experimental replicate, n≥3. (B) Fluorescence intensities from binding curves of fluorescent nanoparticle B1 probe, B1 Streptavidin AlexaFluor™ 647 Conjugate, and B1 Streptavidin APC Conjugate against HHH targets at noted timepoints post fabrication. Data shown are from one experiment. (C) Signal intensity versus concentration of the fluorescent nanoparticle probe, Streptavidin APC Conjugate, and Streptavidin AlexaFluor™ 647 Conjugate. Note that the plateau for the nanoparticles is due to the linear range of the plate reader ending at ∼20,000 AU. Data shown are means ± standard deviation, n=3.

To compare stabilities of fluorescent labels, we evaluated on-target and off-target binding over 16 weeks. The fluorescent nanoparticle B1 probe showed consistent binding affinity for 12 weeks, whereas the Streptavidin APC and Streptavidin AlexaFluor 647 conjugates showed diminished probe affinity at 1 and 2 weeks post fabrication (**Figure 6B**). While binding was diminished, fluorescence intensity over 2 weeks was consistent for all labels, although a slight decrease was observed for Streptavidin APC Conjugate (**Figure S7**). This suggests that the alternative labels have reduced affinity for their targets over time, whereas affinity of the fluorescent nanoparticle probe is maintained for several months.

Finally, we evaluated the relative brightness of the fluorescent nanoparticle and alternative labels. The fluorescent nanoparticles were 4- to 5-fold brighter than Streptavidin SureLight™ APC and Streptavidin AlexaFluor™ 647 Conjugate labels on a molar basis (**Figure 6C**). In summary, the fluorescent nanoparticle probes had superior stability and brightness as compared to alternative labels.

### Use of an alternative nanoparticle core

Finally, we assessed if target specificity was similar when the same functionalization method was applied to a different nanoparticle core, the commercially available Qdot™ 655 ITK™ Carboxyl Quantum Dots. In a plate-based assay, binding affinity for his-tagged Her2 protein was about 2.4-fold higher than background, and the measured EC50 was 55 nM (**Figure 7**). We observed little off-target binding. As noted above, observed affinity can be impacted by both avidity and the detection method, and further studies will be needed to determine why the affinity was lower for the nanoparticle probes with a Qdot core than a FluoSphere core. Overall, this experiment demonstrates that our protocol is robust to swap-in of an alternative core.

**Figure 7.**
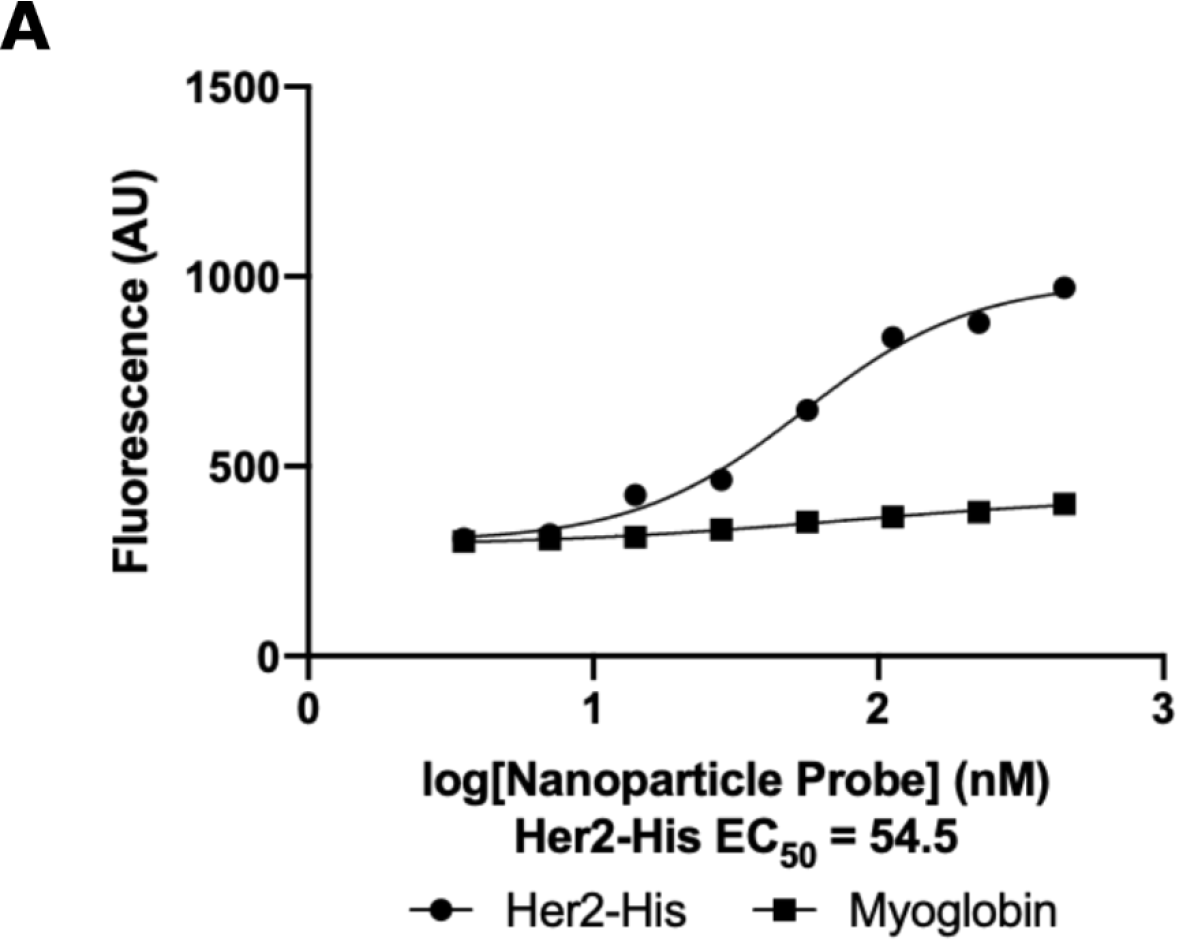
Fluorescent nanoparticle probes can be prepared with a quantum dot core. (A) Binding curves of Qdot fluorescent nanoparticle B1 probe to his-tagged Her2 (on-target) and myoglobin (off-target) proteins. Data show means of duplicates from one experiment.

## Discussion

In these studies, we describe a straightforward method for fabrication of fluorescent nanoparticle probes with the aim of enabling broader use of nanoparticle-based labels. Our protocol utilizes commercially available reagents to create nanoparticle probes with high affinity and brightness, good biocompatibility and colloidal stability, modularity, and long-lasting activity. The first step of our method is nanoparticle passivation. As nanoparticle surfaces tend to non-specifically adsorb proteins^15, 16, 40–43^, dense PEG layers are commonly used to passivate particles due to favorable interactions with water molecules that energetically disfavor interactions with other biomolecules ^35, 42, 44^. For the fluorescent nanoparticle probes presented here, nanoparticles were PEGylated using industry-standard techniques that reduce protein binding^37, 43^. We optimized PEG density and molecular weight to maximize particle passivation while maintaining colloidal stability. As PEG density and molecular weight increased, particle passivation increased. This was expected as short, low-density and low-molecular-weight PEG grafting results in a “mushroom” conformation, ineffective at passivating surfaces, whereas high-density and high-molecular-weight PEG grafting results in “brush” conformations, leading to highly effective passivation^35, 44, 45^ (Supplementary Information, ***PEG Density Calculations***). With optimized PEGylation conditions, off-target nanoparticle probes showed low non-specific binding to proteins. The zeta potential of our nanoparticles was never neutral, however, suggesting that passivation of the particle surface was not complete. We may be able to improve PEGylation of 200-nm particles by increasing the PEG concentration until we see a plateau in zeta potential as we observed for 40 nm particles. Additionally, mixing high and low molecular weight PEGs has been used to improve surface passivation in single molecule detection experiments^46^, and could also improve passivation of nanoparticle surfaces. We could also evaluate other passivating polymer and surfactant combinations^47^.

The next step of our method is the attachment of the conjugation-annealing handle and the DNA aptamer probe. In many probe-labeling protocols, each new probe requires optimization, which can be time-consuming and expensive. Our strategy largely avoids this by achieving modularity via a conjugation-annealing handle, a short DNA sequence designed to anneal aptamer probes on the 5’ end and to covalently attach to the PEG layer through a DBCO moiety on the 3’ end. By utilizing the conjugation-annealing handle, a multitude of probes can be prepared without needing to functionalize each probe with a conjugation handle. Here, we demonstrate this modularity by demonstrating activity of fluorescent nanoparticles functionalized with B1 and its truncated version H3T, both designed to bind his-tags, and an aptamer that binds the VEGF protein^32^. The probe can be attached to the conjugation-annealing handle before or after conjugation to the particle so fluorescent nanoparticles conjugated to the handle can be prepared in large batches. This method could also be used to prepare antibody probes functionalized with a short DNA sequence complementary to the conjugation-annealing handle. Our strategy eliminates problems arising from incompatible chemical modifications of commonly available functionalized nanoparticles and antibodies. One limitation of this approach is that the probes are not covalently attached to the nanoparticles. Although in our hands, nanoparticle probes were stable for months, stronger attachment could be achieved by optimizing the annealing sequences or employing non-natural nucleotides, such as locked nucleic acid^48^.

We used the B1 aptamer in proof-of-principle experiments due to its high affinity and potential utility in other applications. B1 binds his-tags that are used for the isolation of recombinant proteins via immobilized metal affinity chromatography^49^. The fluorescent nanoparticle B1 probe bound both peptides and proteins specifically. Both the B1 aptamer and its truncated version, H3T, bound targets with EC50 values ranging from 3 to 12 nM, lower than the 120 nM affinities previously reported for these aptamers^31^. Although this suggests that attachment to the nanoparticle increases aptamer affinity, it is important to note that we report concentration on a nanoparticle probe basis, but local concentration of probe may be much higher due to the multivalent nature of the nanoparticle probe^20^. Furthermore, the presence of multiple probes per nanoparticle could help stabilize the interaction with the target, decreasing the off-rate and thus reducing the EC50 . This avidity is a major advantage over traditional labels, which only have one probe per fluorescent label. When we compared fluorescent nanoparticle labels to other off-the-shelf labels, we found that the affinity increased as the ratio of probe to label increased. In addition, when we varied the percentage of PEG with a conjugation handle for probe binding, we noted that higher percentages results in increased probe binding to the target. We hypothesize that both observed effects are due to avidity. We note that EC50 measurements are highly dependent on the experimental conditions and do not provide the kinetic information. Studies that utilize approaches like surface plasmon resonance will improve our understanding of how the fluorescent nanoparticle label impacts probe binding kinetics.

The type of nanoparticle used in the creation of fluorescent nanoparticle probes will depend upon the specific research application. The protocol we describe can be used with different nanoparticle types as demonstrated by our validation of Qdot-based probes. B1 probes with a Qdot core showed specific binding to a his-tagged protein and an HHH peptide with EC50 values of about 55 nM. This EC50 is higher than that measured with the FluorSphere-based probe, possibly due to lower affinity, inadequate detection of Qdot fluorescence, or fewer probes per nanoparticle. Thus, this strategy is highly versatile but will likely require some optimization studies when utilizing a new nanoparticle core.

The fluorescent nanoparticle probes described in this work have brightness profiles and stability on par or better than other commercially available labeling techniques, have high affinity for targets, minimal non-specific binding, and are biocompatible. Our nanoparticle probes take inspiration from nanotechnologies used in other applications: Polystyrene nanoparticles have been used *in vitro* to study targeted cellular interactions and uptake by conjugating cell membrane-targeting moieties to the nanoparticle surface^52, 53^ and quantum dots have been functionalized with aptamers for use as theranostic agents in targeted cancer drug delivery applications^54–57^ and as biomolecule sensors^58^. Previous applications have involved use of custom probes, nanoparticles, or conjugation chemistries, making broad adoption challenging. In contrast, each component of the fluorescent nanoparticle probes presented here is commercially available. In addition, our approach is modular, enabling the same core to be used for multiple probes. We successfully generated nanoparticle probes against his-tags and the VEGF protein with little specialized equipment and reagents that were all commercially available. In the future, clinically relevant DNA, RNA, protein, or small-molecule based probes could be conjugated to the particles using a similar technique.

We have highlighted the utility of the nanoparticle probes in studying probe-target interactions in plate-based assays, but this could be extended to single-molecule studies and flow cytometry. The brightness and resistance to photobleaching of single nanoparticle labels make these probes particularly suitable for both applications. In addition, the nanoparticle probes developed in this work could be used for targeted cellular uptake studies, and the surface functionalization techniques could be extended to create nanoparticles capable of controlled release for intracellular biomolecule delivery. Finally, the modularity of the probe attachment approach allows extension of this method to include other types of biological probes such as proteins and small molecules. In sum, the method for fabricating fluorescent nanoparticle probes described could have vast applications with only slight modifications and will enable researchers to achieve fluorescent sensitivity that is unattainable with traditional labeling approaches.

## Materials and Methods

### Materials for nanoparticle functionalization and passivation

Table 1 lists the vendors and catalog numbers for materials used in these studies. Carboxylate- modified FluoSpheres™ and QDot™ nanoparticles were purchased from ThermoFisher. mPEG-amine (molecular weight (MW): 2,000 g/mol) and azide-PEG-amine (MW: 2048 g/mol) were from Creative PEGWorks. DNA conjugation annealing handles and probes were purchased from Integrated DNA Technologies (IDT) or Eurofins Scientific; sequences are listed in Table 3.

**Table 3.**
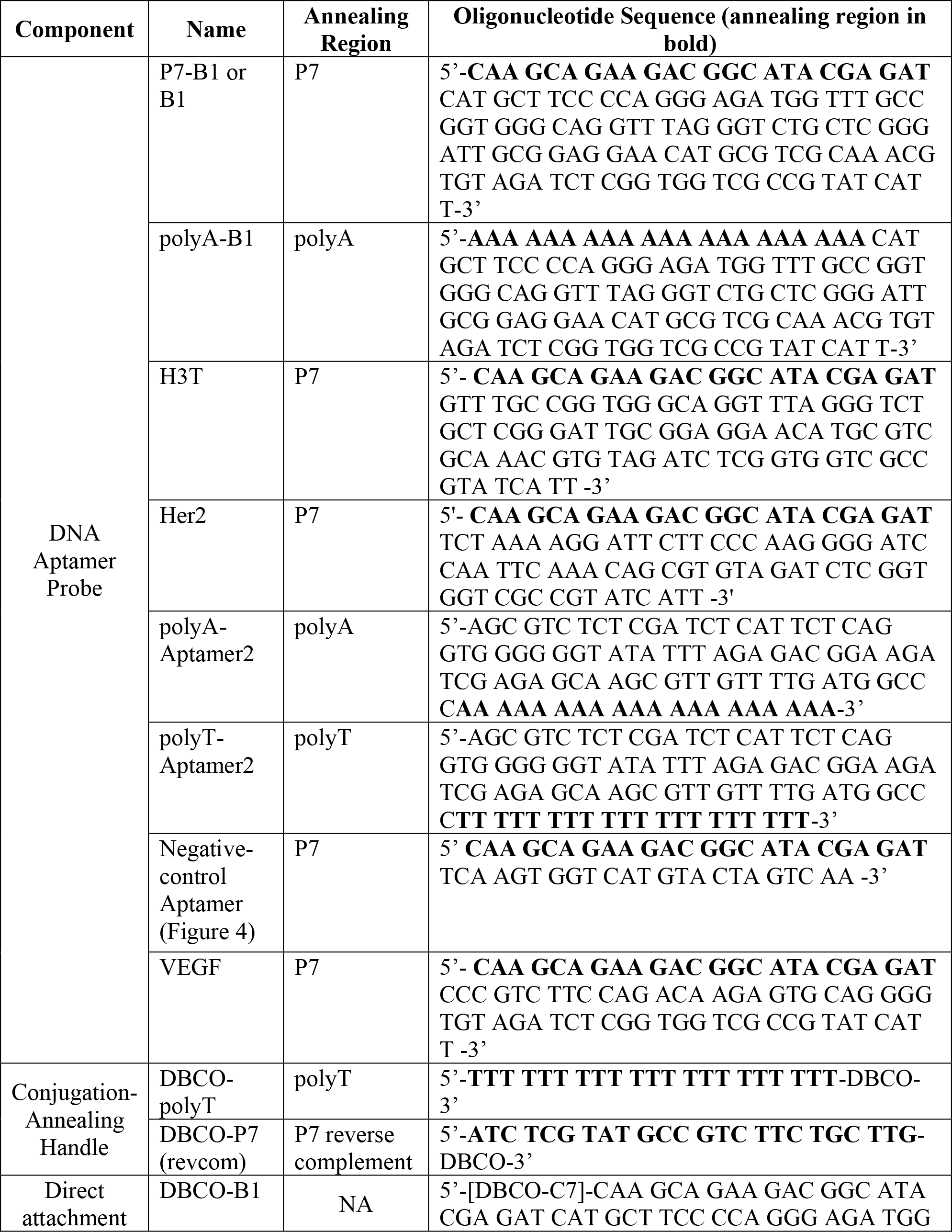
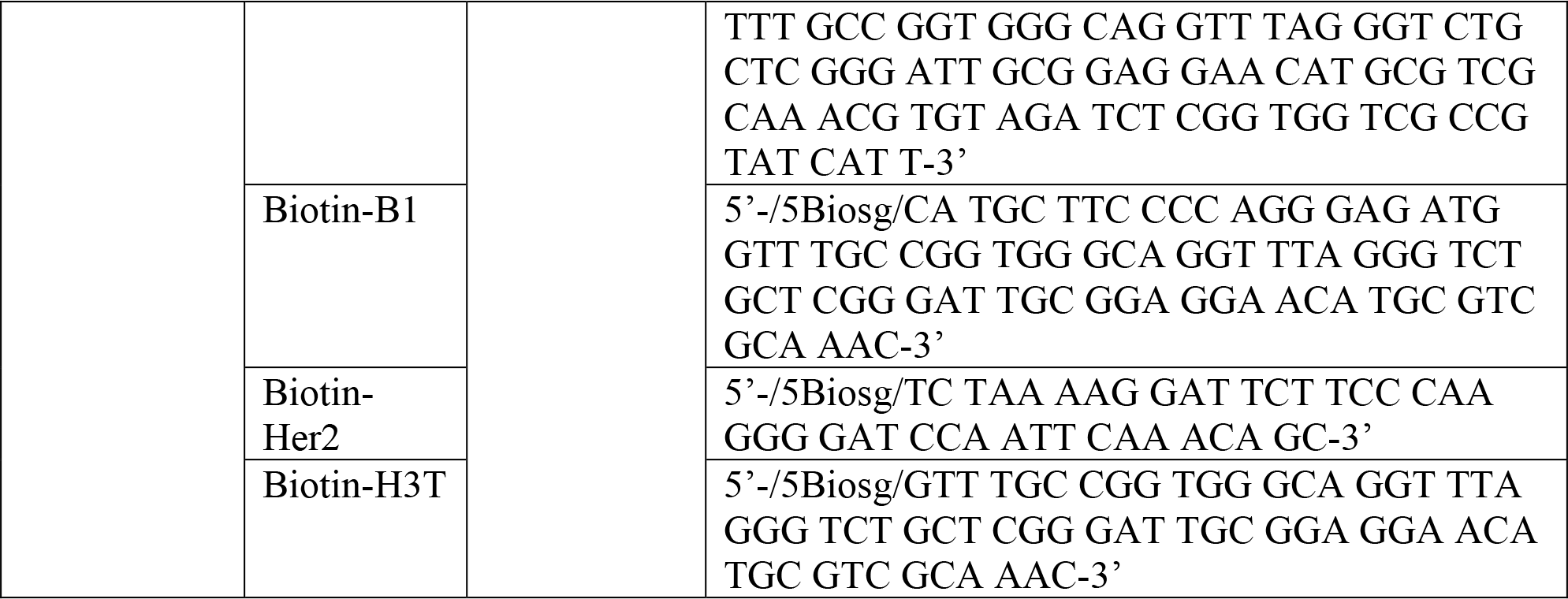
Oligonucleotides for the generation of fluorescent nanoparticle probes

### Nanoparticle dispersion and washing

To ensure that nanoparticles are well-distributed in an aqueous solution, tubes of particles were dispersed by pipetting up and down while partially immersed in a sonication bath (Branson Bransonic Ultrasonic Cleaner 8510R-DTH). Nanoparticle washes were performed as follows: 1) nanoparticles were centrifuged at 31,000 x*g* for 30 minutes (pre-PEGylation) or 60 minutes (post-PEGylation) in a 1.5-mL tube; 2) supernatant was removed, taking care not to disturb the pellet; 3) appropriate volume of buffer was added; and 4) the pellet was redispersed by pipetting up and down while sonicating until no large aggregates of nanoparticles were visible.

### Nanoparticle activation and PEGylation

Carboxylate-modified microspheres were purchased from ThermoFisher. According to ThermoFisher technical support, the carboxylate modified polystyrene particles are generated by copolymerizing a carboxylic acid-containing monomer with styrene. Carboxylate-modified microspheres were activated by reacting 5.3x10^13^ or 1.7x10^11^ nanoparticles/mL for 40-nm and 200-nm nanoparticles, respectively, with 50 mM (N-3-dimethylaminopropyl)-N’- ethylcarbodiimide hydrochloride (EDC; Sigma) and 100 mM N-hydroxysuccinimide sodium salt (NHS; Sigma) in 20 mM 2-(N-morpholino)ethanesulfonic acid (MES; Sigma), pH 6, 500 mM NaCl at 24 °C for 1 hour, shaking at 800 revolutions per minute (RPM) on a ThermoMixer dry block. The protocol outlined here was generally performed on 5.3x10^13^ 40-nm nanoparticles or 2.13x10^10^ 200-nm nanoparticles (see Supplementary Information, ***Fluorescent Nanoparticle Probe Protocol***). Particles were washed, resuspended in 1 mL phosphate buffer saline (PBS; 10 mM Na2HPO4, 1.8 mM KH2PO4, 137 mM NaCl, 2.7 mM KCl, pH 7.4), and washed again.

Particles were resuspended in a 100 mg/mL 95:5 mPEG-amine:azide-PEG-amine solution (Table 1) in PBS such that the PEG:nanoparticle molar ratio was 3.5x10^7^ and 1.4x10^5^ for 200- and 40- nm nanoparticles, respectively. The reaction was incubated at 24 °C for 1 hour shaking at 800 RPM. Then, 250 µl/180 PBS was added to the sample and samples were washed twice with resuspension in 500/125 µl PBS for 40-nm/200-nm nanoparticles, respectively.

Quantum dots were functionalized using the same procedure with the following modifications: the nanoparticles were activated by reacting 7.23x10^14^ nanoparticles/mL with 5 mM EDC and 10 NHS. For PEGylation, nanoparticles were resuspended in 100 mg/mL of a 75:25 mPEG-amine:azide-PEG-amine in PBS to a PEG:nanoparticle molar ratio of 2x10^4^.

### DNA aptamer probe attachment

For both pre-annealing and post-annealing processes, 125 µM of the DNA aptamer probe- complex or the conjugation annealing handle was reacted with particles at a 30,000:1, 125:1, or 25:1 DNA:nanoparticle molar ratio for 200-nm FluoSpheres, 40-nm FluoSpheres, or Qdots, respectively. For pre-annealing, the probe and the handle were combined at concentrations of 125 µM each and incubated at 95 °C for 5 minutes and at room temperature for 10 minutes, and 10x PBS was added to the complex to a final concentration of 1x PBS. The complex was then mixed with PEGylated nanoparticles and reacted at 24 °C overnight with shaking at 800 RPM to allow the DBCO/azide click reaction to proceed. Following incubation, 250/180 µL PBS was added to the sample of 40-nm/200-nM particles, respectively, which was washed twice and resuspended in 500/125 µL PBS for-40 nm/200-nM particles, respectively. The final sample was stored at 4 °C.

For post-annealing, the annealing handle was reacted with 200 nm nanoparticles at 24 °C overnight with shaking at 800 RPM. Nanoparticle were washed twice and resuspended in 180 µL PBS. The DNA aptamer probe was then added to the dry nanoparticle pellet at a concentration of 125 µM and the same molar ratios as the annealing handle, and incubated at 95 °C for 5 minutes and at 24 °C for 10 minutes while shaking at 800 RPM. Nanoparticles were washed twice, resuspended in 125 µl PBS, and stored at 4 °C.

### Preparation of fluorescent probes with alternative off-the-shelf labels

Biotinylated aptamer probes were attached to the following streptavidin conjugates: Streptavidin AlexaFluor™ 647 Conjugate (Thermo Fisher Scientific, S32357), Streptavidin APC Conjugate (ThermoFisher Scientific, S32362), and Streptavidin SureLight™ APC (allophycocyanin, Columbia Biosciences, D3-2212). The conjugation reaction was performed in a buffer containing 10 mM HEPES, 1.2 mM NaCl, 5 mM MgCl2, 5 mM KCl pH 7.4 (HEPES buffer). Streptavidin conjugates at a 1 uM concentration were combined with biotinylated aptamers at a 1:4 molar ratio and incubated for 30 minutes while shaking at 600 RPM in HEPES buffer. Unoccupied biotin binding sites were then blocked by addition of 100 mM D-biotin (Thermo Fisher) in dimethyl sulfoxide (Thermo Fisher) at a 40-molar excess of D-biotin to streptavidin and incubation for 30 minutes while shaking at 600 RPM.

### Fluorescent nanoparticle probe characterization

Dynamic light scattering (DLS) measurements for nanoparticle hydrodynamic diameter, polydispersity index (PDI), and zeta potential were obtained using a Malvern Zetasizer ZSP Zen5600. All reported measurements were carried out in 2% PBS with a 60-second delay between measurements. Aliquots of 2-5 µl of sample were added to 700 µl of 2% PBS and loaded into a cuvette (BrandTech Scientific, 759150). Zeta potential was measured in Folded Capillary Cell cuvettes (Malvern, DTS1070).

qPCR was used to measure the number of probes/particle. The total number of probes/particle was assessed by conducting qPCR once the probes were removed from the particles. Removal was achieved by heating the sample to 95 °C for 10 minutes, centrifuging at 31,000 x*g* for 16 minutes, and collecting supernatant for assessment. Next, qPCR was performed on an Applied Biosystems QuantStudio 3 Real-Time PCR Instrument. A 10-point standard curve with concentrations ranging from 1000 pM to 0.1 pM of the aptamer probe was made in DNase/RNase free water; a no-template control was included. A PCR reaction master mix was prepared with 12.5 µl of 2x PowerUp SYBR Green Master Mix (Applied Biosystems, 100029283), 1.25 µl of 25 µM forward primer, and 1.25 µl of 25 µM reverse primer per sample (see **Table 4** for primer sequences). A 15-µl aliquot of the master mix was combined with 10 µl of sample in a 96-well PCR plate (VWR, 83007-374). The plate was mixed on a Qiagen TissueLyser II for two 25-second intervals at 13 Hz then centrifuged for 2 minutes at 1000 x*g*.

**Table 4.**
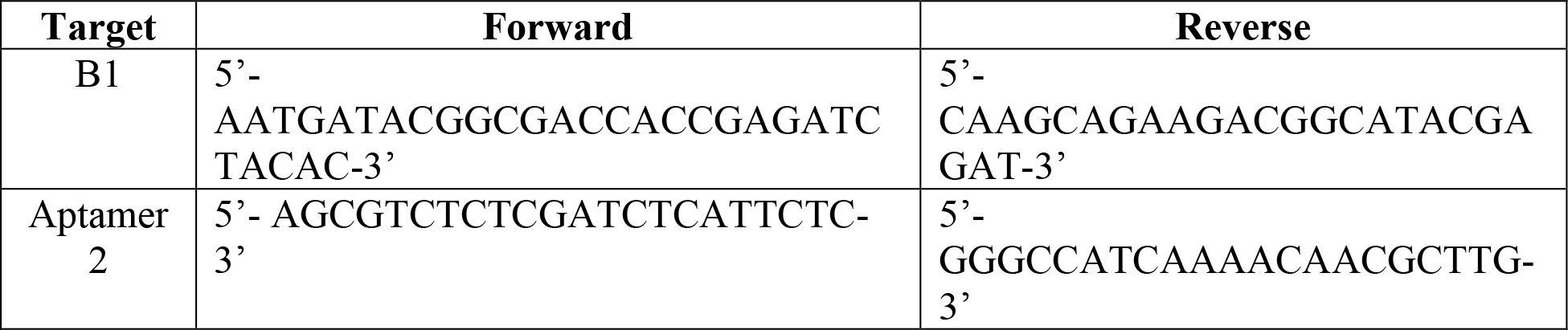
Primers

The qPCR program was as follows: 1) 50 °C for 2 minutes, 2) 95 °C for 2 minutes, 3) 95 °C for 15 seconds, 4) 50 °C for 1 minute, 5) cycle steps 3 and 4 repeated 40 times. The concentration of DNA/nanoparticle was calculated by comparing the Ct values of the experimental sample to the standard curve. Two replicates were run per sample.

### Binding assessment via plate-based assay

The HHH peptide (Cys-biotin-PEG2-Gly-His-His-His-Gly(COOH)) was purchased from Thermo Fisher. It was 84% pure as determined by mass spectrometry; the dominant contaminant was the peptide missing the PEG2 spacer. The PNG peptide was synthesized in-house using an Intavis Multipep RSi synthesizer. Recombinant Her2-his (Sino Biological, 1004H08H) and VEGF (Sino Biological, 11066-HNAH) were resuspended according to the manufacturer’s protocol, aliquoted, and stored at -80 °C. Myoglobin (Sigma-Aldrich, M1882) and *E. coli* lysate (MCLab, ECCL-100) were prepared in 15 mM Na2CO3, 35 mM NaHCO3 (referred to as 50 mM carbonate-bicarbonate buffer), pH 9.6 and stored at 4 °C.

### Peptide ELONA

Wells of streptavidin-coated 384-well plates (Pierce, 15506) were washed twice with 100 µL of wash buffer (0.1% Tween-20, 10 mM HEPES, 1.2 mM NaCl, 5 mM MgCl2, 5 mM KCl, pH 7.4) then incubated with 50 µL of 1 µM biotinylated peptide in peptide binding buffer (1% sheared salmon sperm DNA (Invitrogen, AM9680), 10 mM HEPES, 1.2 mM NaCl, 5 mM MgCl2, 5 mM KCl, pH 7.4) and were shaken at 500 rpm for 30 minutes. Wells were then washed six times with 100 µL wash buffer and blocked with blocking solution (100 µL QBlock (Grace Biolabs, 105106) containing 0.12% Span-80, 0.1% Tween-20, 1% sheared salmon sperm DNA, and 100 µM D-biotin) while shaking at 500 RPM for 30 minutes. A 25-µL aliquot of fluorescently labeled aptamer probe in binding buffer (wash buffer containing 1% sheared salmon sperm DNA and 300 µM dextran sulfate sodium salt (40 kDa, Sigma-Aldrich, 42867)) was added to each well and allowed to incubate for 60 minutes while shaking at 500 RPM. Wells were then washed 6-8 times with 100 µL of wash buffer followed by one wash with 100 µL wash buffer without Tween-20 to remove residual bubbles. Finally, 50 µL of wash buffer without Tween-20 was added to each well, and the plate was read using a Tecan Infinite 200 microplate spectrophotometer.

### Protein ELONAs: Fluorescent nanoparticle probes and Quanta Red HRP Substrate

High-binding, 384-well plates (Corning 3577) were coated with 50 µL of 50 nM protein overnight at 4 °C in 50 mM carbonate-bicarbonate buffer, pH 9.6. The protein solution was washed out with 100 µL of wash buffer per well and blocked with 100 µL of blocking solution for 90 minutes. The plates were washed six times with 100 µL of wash buffer per well and then 25 µL biotinylated aptamer probes or 25 µL of fluorescent nanoparticle probes were added in binding buffer and incubated for 1 hour. For biotinylated probes, wells were then washed three times with wash buffer, and 50 µL of binding buffer containing 5 µg/mL Streptavidin Poly-HRP (ThermoFisher Scientific 21140) was added to each well and incubated for 30 minutes with shaking at 500 RPM. A solution of QuantaRed HRP substrate (ThermoFisher Scientific, 15159) was prepared according to manufacturer’s protocol, and 50 µL of the substrate was added to each well and incubated for 5 minutes followed by addition of 5 µL stop solution. For fluorescent nanoparticles probes, wells were washed six times with wash buffer after the 1 hour incubation with probes. Fluorescence was measured on a Tecan Infinite 200 microplate spectrophotometer. *Determination of EC50 values*

After obtaining raw fluorescence intensities from ELONAs, the concentration of nanoparticles used was converted to logarithmic scale. GraphPad Prism was used to perform the half maximal effective concentration (EC50) analysis. The figures were analyzed using the “Sigmoidal, 4PL, X is log(concentration)” option. This fits a four-parameter logistic curve (4PL) and the Hill Slope from the data. Then, EC50 was obtained from the interpolated “Best-fit values” post analysis.

### Comparison to other fluorescent labels

Biotinylated aptamer probes conjugated to alternative off-the-shelf labels were stored at concentrations of 1 µM in 10 mM HEPES, 1.2 mM NaCl, 5 mM MgCl2, 5 mM KCl, pH 7.4 at 4 °C. Stability of these alternative fluorescent probes was determined by conducting a peptide ELONA immediately and 1-3 weeks post-conjugation. Fluorescent nanoparticle B1 probes were stored in 500 µl PBS at 4 °C. Stability of the fluorescent nanoparticle probes was assessed by performing a peptide ELONA immediately and 8, 12, and 16 weeks post-conjugation using a 3- fold serial dilution of probe from 100 nM to 7 fM in 100 µl 1X NV buffer using a 384-well black microplate (Corning 3601).

### Data Availability

The datasets generated during and/or analyzed during the current study are available from the corresponding author on request.

## Acknowledgements

We wish to acknowledge Elliott SoRelle, Tim Ruckh, Ming Tan, David Stern, Dmitri Gremyachinskiy, Tural Aksel, and the scientists at Malvern Panalytical for their advice and consultation in this project.

## Author Contributions

TER, PM, and CMS wrote the main manuscript text, and CMS, MB, and TER prepared figures. TER, CMS, MB, SST, RPG, JKR, and PM contributed to the planning, experimental design, and data analysis. CMS, MB, TER, DA, and JKR contributed to experimental execution and data compilation. All authors reviewed the manuscript.

## Additional Information

The authors declare no competing interests.

## Notes

### Competing Interest Statement

The authors have declared no competing interest.

## References

1. Ueno, T. & Nagano, T. Fluorescent probes for sensing and imaging. Nat. Methods 8, 642–645 (2011).

2. Goldshtein, H., Hausmann, M. J. & Douvdevani, A. A rapid direct fluorescent assay for cell-free DNA quantification in biological fluids. Ann. Clin. Biochem. Int. J. Lab. Med. 46, 488–494 (2009).

3. Kapanidis, A. N. & Weiss, S. Fluorescent probes and bioconjugation chemistries for single-molecule fluorescence analysis of biomolecules. J. Chem. Phys. 117, 10953–10964 (2002).

4. Lo Giudice, M. C., Herda, L. M., Polo, E. & Dawson, K. A. In situ characterization of nanoparticle biomolecular interactions in complex biological media by flow cytometry. Nat. Commun. 7, 13475 (2016).

5. Yameen, B. et al. Insight into nanoparticle cellular uptake and intracellular targeting. J. Control. Release 190, 485–499 (2014).

6. Welsher, K. & Yang, H. Multi-resolution 3D visualization of the early stages of cellular uptake of peptide-coated nanoparticles. Nat. Nanotechnol. 9, 198–203 (2014).

7. Wang, T. et al. Size-Dependent Regulation of Intracellular Trafficking of Polystyrene Nanoparticle-Based Drug-Delivery Systems. ACS Appl. Mater. Interfaces 9, 18619–18625 (2017).

8. Shi, D. et al. Fluorescent Polystyrene-Fe 3 O 4 Composite Nanospheres for In Vivo Imaging and Hyperthermia. Adv. Mater. 21, 2170–2173 (2009).

9. Cheng, L. et al. Highly-sensitive multiplexed in vivo imaging using pegylated upconversion nanoparticles. Nano Res. 3, 722–732 (2010).

10. Ma, D.-L., He, H.-Z., Leung, K.-H., Chan, D. S.-H. & Leung, C.-H. Bioactive Luminescent Transition-Metal Complexes for Biomedical Applications. Angew. Chemie Int. Ed. 52, 7666–7682 (2013).

11. Chudakov, D. M., Matz, M. V., Lukyanov, S. & Lukyanov, K. A. Fluorescent Proteins and Their Applications in Imaging Living Cells and Tissues. Physiol. Rev. 90, 1103–1163 (2010).

12. Resch-Genger, U., Grabolle, M., Cavaliere-Jaricot, S., Nitschke, R. & Nann, T. Quantum dots versus organic dyes as fluorescent labels. Nat. Methods 5, 763–775 (2008).

13. Buck, S. et al. Nanoscale probes encapsulated by biologically localized embedding (PEBBLEs) for ion sensing and imaging in live cells. Talanta 63, 41–59 (2004).

14. Petrizza, L. et al. Dye-doped silica nanoparticle probes for fluorescence lifetime imaging of reductive environments in living cells. RSC Adv. 6, 104164–104172 (2016).

15. Giepmans, B. N. G., Adams, S. R., Ellisman, M. H. & Tsien, R. Y. The Fluorescent Toolbox for Assessing Protein Location and Function. Science (80-.). 312, 217–224 (2006).

16. Reisch, A. & Klymchenko, A. S. Fluorescent Polymer Nanoparticles Based on Dyes: Seeking Brighter Tools for Bioimaging. Small 12, 1968–1992 (2016).

17. Gref, R. et al. ‘Stealth’ corona-core nanoparticles surface modified by polyethylene glycol (PEG): influences of the corona (PEG chain length and surface density) and of the core composition on phagocytic uptake and plasma protein adsorption. Colloids Surfaces B Biointerfaces 18, 301–313 (2000).

18. Coto-García, A. M. et al. Nanoparticles as fluorescent labels for optical imaging and sensing in genomics and proteomics. Anal. Bioanal. Chem. 399, 29–42 (2011).

19. Larson, D. R. et al. Silica Nanoparticle Architecture Determines Radiative Properties of Encapsulated Fluorophores. Chem. Mater. 20, 2677–2684 (2008).

20. Pinaud, F. et al. Advances in fluorescence imaging with quantum dot bio-probes. Biomaterials 27, 1679–1687 (2006).

21. Alivisatos, A. P., Gu, W. & Larabell, C. Quantum Dots as Cellular Probes. Annu. Rev. Biomed. Eng. 7, 55–76 (2005).

22. Hartlen, K. D., Athanasopoulos, A. P. T. & Kitaev, V. Facile preparation of highly monodisperse small silica spheres (15 to >200 nm) suitable for colloidal templating and formation of ordered arrays. Langmuir 24, 1714–1720 (2008).

23. Valizadeh, A. et al. Quantum dots: synthesis, bioapplications, and toxicity. Nanoscale Res. Lett. 7, 480 (2012).

24. Nightingale, A. M. & de Mello, J. C. Microscale synthesis of quantum dots. J. Mater. Chem. 20, 8454 (2010).

25. LaBauve, A. E. et al. Lipid-Coated Mesoporous Silica Nanoparticles for the Delivery of the ML336 Antiviral to Inhibit Encephalitic Alphavirus Infection. Sci. Rep. 8, 13990 (2018).

26. Kumar, R. et al. Covalently Dye-Linked, Surface-Controlled, and Bioconjugated Organically Modified Silica Nanoparticles as Targeted Probes for Optical Imaging. ACS Nano 2, 449–456 (2008).

27. Wu, C. et al. Bioconjugation of Ultrabright Semiconducting Polymer Dots for Specific Cellular Targeting. J. Am. Chem. Soc. 132, 15410–15417 (2010).

28. Tang, B., Tang, B., Cheang, Wang, S. & Pro Xu. Promising plasmid DNA vector based on APTES-modified silica nanoparticles. Int. J. Nanomedicine 7, 1061 (2012).

29. Kumar, R. et al. Modified Silica Nanoparticles as Targeted Probes for Optical Imaging. 2, 449– 456 (2008).

30. Rosenthal, S. J., Chang, J. C., Kovtun, O., McBride, J. R. & Tomlinson, I. D. Biocompatible Quantum Dots for Biological Applications. Chem. Biol. 18, 10–24 (2011).

31. Bartnicki, F., Kowalska, E., Pels, K. & Strzalka, W. Imidazole-free purification of His 3 -tagged recombinant proteins using ssDNA aptamer-based affinity chromatography. J. Chromatogr. A 1418, 130–139 (2015).

32. Potty, A. S. R. et al. Biophysical characterization of DNA aptamer interactions with vascular endothelial growth factor. Biopolymers 91, 145–156 (2009).

33. Smith, M. E. B. et al. US Patent 8,563,477 B2 Modified Molecular Arrays. (2013).

34. Bhattacharjee, S. DLS and zeta potential – What they are and what they are not? J. Control. Release 235, 337–351 (2016).

35. Yang, Q. et al. Evading immune cell uptake and clearance requires PEG grafting at densities substantially exceeding the minimum for brush conformation. Mol. Pharm. 11, 1250–1258 (2014).

36. Hoo, C. M., Starostin, N., West, P. & Mecartney, M. L. A comparison of atomic force microscopy (AFM) and dynamic light scattering (DLS) methods to characterize nanoparticle size distributions. J. Nanoparticle Res. 10, 89–96 (2008).

37. Suk, J. S., Xu, Q., Kim, N., Hanes, J. & Ensign, L. M. PEGylation as a strategy for improving nanoparticle-based drug and gene delivery. Adv. Drug Deliv. Rev. 99, 28–51 (2016).

38. Kim, E.-Y. et al. A real-time PCR-based method for determining the surface coverage of thiol-capped oligonucleotides bound onto gold nanoparticles. Nucleic Acids Res. 34, e54–e54 (2006).

39. Gijs, M. et al. Improved aptamers for the diagnosis and potential treatment of HER2-positive cancer. Pharmaceuticals 9, 15–19 (2016).

40. Amoozgar, Z. & Yeo, Y. Recent advances in stealth coating of nanoparticle drug delivery systems. WIREs Nanomedicine and Nanobiotechnology 4, 219–233 (2012).

41. Verma, A. & Stellacci, F. Effect of Surface Properties on Nanoparticle-Cell Interactions. Small 6, 12–21 (2010).

42. Perry, J. L. et al. PEGylated PRINT Nanoparticles: The Impact of PEG Density on Protein Binding, Macrophage Association, Biodistribution, and Pharmacokinetics. Nano Lett. 12, 5304– 5310 (2012).

43. Otsuka, H., Nagasaki, Y. & Kataoka, K. PEGylated nanoparticles for biological and pharmaceutical applications. Adv. Drug Deliv. Rev. 64, 246–255 (2012).

44. Damodaran, V. B., Fee, C. J., Ruckh, T. & Popat, K. C. Conformational Studies of Covalently Grafted Poly (ethylene glycol) on Modified Solid Matrices Using X-ray Photoelectron Spectroscopy. 26, 7299–7306 (2010).

45. Perry, J. L., et al. PEGylated PRINT Nanoparticles: The Impact of PEG Density on Protein Binding, Macrophage Association, Biodistribution, and Pharmacokinetics. (2012).

46. Chandradoss, S. D. et al. Surface Passivation for Single-molecule Protein Studies. J. Vis. Exp. 1–8 (2014) doi:10.3791/50549.

47. Hadjesfandiari, N. & Parambath, A. Stealth coatings for nanoparticles. in Engineering of Biomaterials for Drug Delivery Systems 345–361 (Elsevier, 2018). doi:10.1016/B978-0-08-101750-0.00013-1.

48. Hagedorn, P. H. et al. Locked nucleic acid: modality, diversity, and drug discovery. Drug Discov. Today 23, 101–114 (2018).

49. Lichty, J. J., Malecki, J. L., Agnew, H. D., Michelson-Horowitz, D. J. & Tan, S. Comparison of affinity tags for protein purification. Protein Expr. Purif. 41, 98–105 (2005).

50. Hong, S. et al. The Binding Avidity of a Nanoparticle-Based Multivalent Targeted Drug Delivery Platform. Chem. Biol. 14, 107–115 (2007).

51. Vauquelin, G. & Charlton, S. J. Exploring avidity: understanding the potential gains in functional affinity and target residence time of bivalent and heterobivalent ligands. Br. J. Pharmacol. 168, 1771–1785 (2013).

52. Lin, A. et al. Shear-regulated uptake of nanoparticles by endothelial cells and development of endothelial-targeting nanoparticles. J. Biomed. Mater. Res. Part A 93, 833–842 (2010).

53. Braeckmans, K. et al. Transport of nanoparticles in cystic fibrosis sputum and bacterial biofilms by single-particle tracking microscopy. Nanomedicine 8, 935–949 (2012).

54. Bagalkot, V. et al. Quantum dot - Aptamer conjugates for synchronous cancer imaging, therapy, and sensing of drug delivery based on Bi-fluorescence resonance energy transfer. Nano Lett. 7, 3065–3070 (2007).

55. Li, Z. et al. Aptamer-conjugated dendrimer-modified quantum dots for cancer cell targeting and imaging. Mater. Lett. 64, 375–378 (2010).

56. Liu, H., Xu, S., He, Z., Deng, A. & Zhu, J. Supersandwich Cytosensor for Selective and Ultrasensitive Detection of Cancer Cells Using Aptamer-DNA Concatamer-Quantum Dots Probes. Anal. Chem. 85, 1–5 (2013).

57. Chen, X. C. et al. Quantum dot-labeled aptamer nanoprobes specifically targeting glioma cells. Nanotechnology 19, (2008).

58. Kim, G. Il, Kim, K. W., Oh, M. K. & Sung, Y. M. The detection of platelet derived growth factor using decoupling of quencher-oligonucleotide from aptamer/quantum dot bioconjugates. Nanotechnology 20, (2009).

