## Supplemental Information for "Modular Fluorescent Nanoparticle DNA Probes for Detection of Peptides and Proteins"

*To whom correspondence should be addressed

### ***Supplemental Results and Discussion***

*Determination of PEG density for 40-nm particles*

To evaluate how PEG density impacted passivation of 40-nm particles, we varied the PEG:nanoparticle ratio and measured the resulting surface charge neutralization using zeta potential measurements. Based on theoretical models, we reduced the PEG:nanoparticle ratio to approximately $10^5$ PEG molecules per nanoparticle and tested several concentrations within that order of magnitude. These studies indicated that there was similar surface charge of -10 mV for each experimental group (**Figure S1**). From this, we concluded that $1.4 \times 10^5$ PEG:nanoparticle was sufficient for passivating 40-nm nanoparticles.

*Binding assessment of varied functional PEG ratios*

To evaluate the ratio of functionalized to non-functionalized PEG required for probe attachment and successful detection, we varied the ratio of amine-PEG-azide to mPEG-amine during nanoparticle fabrication. Fluorescent nanoparticle probes with ratios of mPEG-amine:amine-PEG-azide of 75:25, 95:5, 99.5:0.5, 99.95:0.05, and 99.995:0.005 were fabricated, conjugated to B1 probes, and evaluated in plate-based binding studies. The ratio of 75:25 and 95:5 showed similar binding, while the lower ratios showed a decline in on-target binding that correlated with amine-PEG-azide density (**Figure S2**). As the amine-PEG-azide contains the reactive group for probe attachment, it is expected that if the DNA aptamer probe-complex is in excess of the amine-PEG-azide, the number of probes per particle will increase with the amine-PEG-azide fraction and thus increase on-target binding. This trend was observed for the ratios of 95:5 and lower. However, it is possible that at higher amine-PEG-azide densities, the aptamer-probe complex is no longer in excess or steric hinderance between adjacent probes impacts probe-target interactions. We hypothesize that one of these two mechanisms was responsible for the only marginal increase in binding between the 95:5 and 75:25 groups. For these proof-of-principle studies, we utilized the 95:5 mPEG-amine:amine-PEG-azide ratio for PEGylation of the fluorescent nanoparticle probes to maximize binding efficiency while minimizing cost.

*Characterization of the conjugation-annealing handle conjugation*

To validate the conjugation of the conjugation-annealing handle to the PEG-azide layer of the fluorescent nanoparticles, we quantified the attachment of a fluorescently-labeled conjugation-annealing handle. Using our standard nanoparticle fabrication process, we reacted particles with 125 µM (standard), 12.5 µM, 1.25 µM, and 0 µM of a conjugation-annealing handle that contained a Cy5 dye (“DBCO-Oligo-Cy5”). As a control, nanoparticles were also reacted with 12.5 µM of a conjugation-annealing handle that contained a Cy5 dye but no DBCO functional group (“Oligo-Cy5”). A standard plate reader was used to correlate nanoparticle number with fluorescently labeled oligonucleotide number. Nanoparticles reacted with 0 µM DBCO-Oligo-Cy5 and 12.5 µM Oligo-Cy5 resulted in less than 1 oligonucleotide/nanoparticle, while nanoparticles reacted with 1.25, 12.5, and 125 µM DBCO-oligo-Cy5 resulted in about 1.5, 2.7, and 21oligonucleotides/particle, respectively (**Supplementary Figure S3**). These results indicate that the attachment of the conjugation-annealing handle to the particles is specific and can be modulated through input concentration. In addition, the ratio of ~20 oligos per 40 nm particle is similar to the ~400 oligos per 200 nm particle observed in qPCR (**Figure 3B**) if surface area differences are accounted for. Overall, these results indicate that the conjugation-annealing handle attaches to the particle specifically and the density can be controlled.

*Pre-anneal versus post-anneal probe attachment*

We evaluated two different approaches for probe attachment to the particle: the "pre-anneal" and "post-anneal" approaches. In pre-annealing, the probe is attached to the fluorescent nanoparticle B1 probe by pre-annealing of conjugation-annealing handle to DNA aptamer probe prior to attachment to the particle. In post-annealing, the conjugation-annealing handle is conjugated to the particle and then the DNA aptamer probe is annealed. The pre-annealing and post-annealing methods were compared through assessment of B1 binding to his-tagged Her2 protein (on-target) and myoglobin (off-target). Pre-annealed probes showed higher binding than the post-annealing approach to on-target proteins (5.5-fold and 2.5-fold binding over background, respectively (**Figure S5**). Little non-specific binding was observed irrespective of the attachment strategy. It may be possible to optimize the post-annealing approach in studies beyond the scope of this proof-of-principle analysis. First, the concentration of probe could be increased to ensure attachment to all available conjugation-annealing handles. Second, the annealing handle could include a spacer region to reduce steric hinderance due to the PEG layer. Finally, the annealing region could be extended to increase the likelihood of achieving annealing. For the proof-of-principle studies described in this work, the pre-annealing method was used.

*Binding assessment of alternative conjugation strategies*

As our probe annealing attachment technique is unique, we compared on-target binding by probe prepared using our pre-annealing attachment approach to a more standard direct probe conjugation attachment. In the direct conjugation strategy, the 5'-DBCO functional group on the DNA probe was conjugated to the PEG-azide. The original design, using the annealing approach for probe attachment, showed higher binding to its target than the direct DBCO conjugation strategy (4.3-fold versus 2.4-fold binding over background, respectively; **Figure S6**). This suggest that our annealing approach works as well or better than more standard approaches.

*Fluorescent stability of commercially available probes*

Our fluorescent nanoparticle probes bound with lower $EC_{50}$ values than commercially available labels (**Figure 6**). To investigate if this was due to changes in binding affinity or reduction in fluorescent signal, the brightness of probes was assessed immediately after preparation and at 1 and 2 weeks. Minimal change in fluorescent intensities was observed for the commercially available labels over the course of two weeks (**Supplementary Figure S7**). This study suggests that the reduction in binding signal observed in the commercially available labels is due to loss of binding affinity, not loss in fluorescent intensity. This provides additional motivation to use the fluorescent nanoparticle probes, as they are more stable.

*Relationship between fluorescent intensity and label concentration*

The linear range for yellow-green nanoparticles, dark-red nanoparticles, Streptavidin-647 and Streptavidin-APC was determined for the fluorescent plate reader used in binding assay. The linear range was determined to be ~50-18,000 AU for 488/535 and ~100-10,000 for 635/680 em/ex (**Supplementary Figure S12**). Unless indicated in the figure caption, all data generated in these studies fell within the linear range of the plate reader.

*Experimental data replicates*

**Supplementary Figures S8-11** show additional replicates of the main text figures.

***Supplemental Methods***

*PEG density determination for 40-nm particles*

PEGylation was carried out as described in the main text Materials and Methods section "Nanoparticle activation and PEGylation". Particles were resuspended in a 100 mg/mL of a 95:5

mPEG-amine:azide-PEG-amine (MW of PEGs was 2000 g/mol) in PBS at PEG:nanoparticle molar ratios of $1.4x10^5$, $2.1x10^5$, and $2.8x10^5$. Additional sets of 40-nm particles were resuspended in 100 mg/mL of a 75:25, 99.5:0.5, 99.95:0.05, or 99.995:0.005 mPEG-amine:azide-PEG-amine ratios. The reactions were incubated at 24 °C with shaking at 800 RPM on a ThermoMixer dry block. After 1 hour, 250 µl PBS was added, samples were washed twice, and resuspended in 500 µl PBS. Zeta potential measurements were obtained as described in the main text Materials and Methods section "Fluorescent nanoparticle probe characterization".

*Characterization of conjugation-annealing handle conjugation*

Nanoparticle functionalization was carried out according to the method described through PEGylation. After PEGylation, nanoparticles were reacted with 125 µM (standard), 12.5 µM, 1.25 µM, and 0 µM of a conjugation-annealing handle that contained a Cy5 dye ("DBCO-Oligo-Cy5"; /5DBCOTEG/TGTGGAGAGGAAGATGGTA/3Cy5Sp/). As a control, nanoparticles were also reacted with 12.5 µM of a conjugation-annealing handle that contained a Cy5 dye but no DBCO functional group ("Oligo-Cy5"; TGTGGAGAGGAAGATGGTA/3Cy5Sp/). The remainder of the standard protocol was followed. The fluorescent intensities of nanoparticles were measured on a Tecan Spark at em/ex of 488/535 (nanoparticles) and 610/670 (Cy5) in a black 96-well plate. A standard curve of nanoparticles and the fluorescent oligonucleotides was used to determine the number of oligos/nanoparticle.

*Alternative probe conjugation strategies*

Oligonucleotides were purchased from Integrated DNA Technologies (IDT) for DNA attachment through a conjugation-annealing handle or from Eurofins Scientific for the direct conjugation of the DBCO-B1 probe (**Table 1**). The attachment of DNA was achieved by pre- or post-annealing the probe to the particle, as described in the main text Material and Methods section "DNA aptamer probe attachment" (**Figure 1A**). Direct conjugation of the probe to the PEG layer was performed as a comparison to more standard approaches. For direct conjugation, 40-nm nanoparticles were PEGylated as described in the main text. The DBCO-B1 probe was reacted with azide-PEGylated particles at a 300:1 ratio in PBS at 24 °C overnight with shaking at 800 RPM. Following incubation, 180 µl PBS was added, and samples were washed twice and resuspended in 500 µl PBS. The final sample was stored at 4 °C.

*In-house peptide synthesis*

Peptides were synthesized on an Intavis Multipep RSi synthesizer. All fluorenylmethoxycarbonyl protected amino acids (Fmoc-amino acids) and 1-[bis(dimethylamino)methylene]-1H-1,2,3-triazolo[4,5-b]pyridinium 3-oxide hexafluorophosphate (HATU) were purchased from AAPPTec. All other solvents and reagents were purchased from Sigma or ACROS. For 2-µmole-scale synthesis on Rink Amide AM resin (200-400 mesh), 0.5 M Fmoc-amino acid solutions in anhydrous dimethylformamide (DMF) were activated by 0.5 M HATU. Activated Fmoc-amino acids were coupled to amine group on N-terminus of the previously added amino acid (or amine group of Rink amide resin in case of first amino acid coupling) using 4.0 N *N*-methylmorpholine. The Fmoc group was removed by incubation in 20% piperidine in DMF prior to addition of next incoming amino acid. The first four cycles of peptide synthesis were double coupled for 15 and 25 minutes followed by double coupling for 20 and 30 minutes for remaining cycles. Deprotection and cleavage of peptides from resin were achieved in 94% trifluoroacetic acid (TFA), 2.5% deionized water, 2.5% 1,2-ethane-di-thiol, 1% triisopropylsilane. Peptides were precipitated by adding tert-butyl methyl ether to concentrated peptidyl-TFA solution. The precipitates were dried in a CentriVap, suspended in 500 µL water, and purified using Waters Sep-Pak C-18 plates[1]. Peptides were identified by their m/z value on AB Sciex MALDI MS at the Canary Center, Stanford University.

*Fluorescent stability studies of alternative labels*

Biotinylated aptamer probes conjugated to alternative off-the-shelf labels prepared as described in main text Materials and Methods section “Preparation of fluorescent probes with alternative off-the-shelf labels” were stored at concentrations of 1 µM in 10 mM HEPES, 1.2 mM NaCl, 5 mM $MgCl_2$, 5 mM KCl, pH 7b.4 at 4 °C. The stability of the conjugated fluorescent entity was determined by performing a 3-fold serial dilution from 300 nM to 137 pM in 100 µL 1X NV buffer in a 96-well black microplate (Corning). The fluorescence intensity of each well was used to generate a linear curve of the fluorescent probe immediately and at 1- and 2-weeks post-conjugation.

*Limit of detection for fluorescent plate reader*

For each label used in these studies, a titration with two-fold dilutions into PBS was performed in a 96-well black plate. The fluorescent intensities were measured as described in the main text and graphed in Excel. The linear region for each label was determined evaluating the quality of fit after fitting a regression line using the method of least squares in Excel Software.

***Fluorescent Nanoparticle Probe Protocol***

<u>Buffers:</u>
Reaction buffer: 20 mM MES, 500 mM NaCl, pH 6
Wash buffer: PBS; 10 mM $Na_2HPO_4$, 1.8 mM $KH_2PO_4$, 137 mM NaCl, 2.7 mM KCl, pH 7.4

<u>Nanoparticle Wash:</u>
*Nanoparticle washes are carried out throughout this protocol and consist of the following steps:*
1. Centrifuge nanoparticles at 31,000 x*g* for 30 minutes prior to PEGylation or 60 minutes post PEGylation in a 1.5-mL tube to form a pellet.
2. Remove supernatant, taking care not to disturb the pellet.
3. Add appropriate buffer and volume to the pellet, as noted in procedure.

4. Redisperse the pellet by pipetting up and down while the nanoparticle tube is partially immersed in a standard laboratory sonication bath (Branson Bransonic Ultrasonic Cleaner 8510R-DTH) until no large aggregates of nanoparticles are visible (usually 10-30 seconds).

Procedure:

*This procedure is written for any scale of nanoparticle preparation, but suggested masses, volumes, and concentrations are listed for 40-nm and 200-nm particle preparations used throughout the protocol.*

1. Redisperse the stock tube of carboxylate-modified microspheres (see **Table 1** for more information) for 10 seconds following step 4 of "Nanoparticle Wash" to ensure nanoparticles are well distributed in solution.
2. Pipette $5.3 \times 10^{13}$ 40-nm nanoparticles or $2.13 \times 10^{10}$ 200-nm nanoparticles into 1.5-mL Eppendorf tubes. (Note: Pellets do not form well in 2 mL tubes.) Suggested volumes:
    a. 40-nm particles: 60 µL ($5.3 \times 10^{13}$ nanoparticles) from a stock of $8.8 \times 10^{14}$ particles/mL.
    b. 200-nm particles: 6.25 µL ($2.13 \times 10^{10}$ nanoparticles) from a stock of $3.4 \times 10^{12}$ particles/mL.
3. Activate nanoparticles with EDC/NHS at $5.3 \times 10^{13}$ 40-nm nanoparticles/mL or $1.7 \times 10^{11}$ 200-nm nanoparticles/mL in a solution of 50 mM EDC and 100 mM NHS in reaction buffer at 24 °C for 1 hour with shaking at 800 revolutions per minute (RPM) on a ThermoMixer dry block. Dissolve EDC and NHS directly into reaction buffer immediately before use and carry out steps 4 and 5 quickly to reduce NHS hydrolysis. Suggested volumes:
    a. 40-nm particles: Add 500 µL of 100 mM EDC and 200 mM NHS (both dissolved in reaction buffer) to 60 µL of nanoparticle stock.
    b. 200-nm particles: Add 62.5 µL of 100 mM EDC and 200 mM NHS (both dissolved in reaction buffer) to 60 µL of nanoparticle stock.
4. Wash particle pellet and resuspend in 1 mL wash buffer.
5. Wash particles and redisperse in PEG solution:
    a. Dissolve mPEG-amine and azide-PEG-amine (see **Table 1** for details) at 100 mg/mL concentration in wash buffer.
    b. Resuspend particles in 95:5 volume ratio of mPEG-amine:azide-PEG-amine such that PEG:nanoparticle molecular ratio is $3.5 \times 10^{7}$ and $1.4 \times 10^{5}$ for 200- and 40-nm nanoparticles, respectively. Suggested volumes:
        i. 40-nm particles: 235 µL 100 mg/mL mPEG-amine and 12.7 µL 100 mg/mL azide-PEG-amine.
        ii. 200-nm particles: 23.5 µL 100 mg/mL mPEG-amine and 1.3 µL 100 mg/mL azide-PEG-amine.
6. Incubate at 24 °C for 1 hour with shaking at 800 RPM on a ThermoMixer dry block. (Note: This incubation step may be allowed to proceed overnight.)
7. Add 1 µL wash buffer/µL PEG solution for 40-nm particles and 7 µL wash buffer/µL PEG solution for 200-nm particles. Suggested volumes:
    a. 40-nm particles: 250 µL wash buffer.
    b. 200-nm particles: 180 µL wash buffer.
8. Wash and redisperse particles in 2 µL wash buffer/µL PEG solution PBS for 40-nm particles and 5 µL wash buffer/µL PEG solution for 200-nm particles. (Note: The pellet will be less defined, and some nanoparticles may remain in solution after this step.) Suggested volumes:

    a. 40-nm particles: 500 µL wash buffer.
    b. 200-nm particles: 125 µL wash buffer.
9. Combine particles with the conjugation-annealing handle or probe for the pre-anneal or post-anneal probe attachment approach. React 125 µM of the DNA aptamer probe-complex or the conjugation annealing handle with particles at a 30,000:1 or 125:1 DNA:nanoparticle molar ratio for 200- and 40-nm particles, respectively:
    a. Pre-anneal:
        i. Pre-anneal the conjugation-annealing handle and probe (see **Table 3** for details) by combining 125 µM conjugation annealing handle and 125 µM DNA aptamer probe in equal volumes and incubate at 95 °C for 5 minutes. Centrifuge briefly to remove condensation from the tube lid. Incubate on benchtop for 10-15 minutes to allow the two complementary regions of DNA to anneal. Suggested volumes:
            1. 40-nm particles: Combine 88 µl each of 125 µM stocks of conjugation annealing handle and probe.
            2. 200-nm particles: Combine 8.5 µl each of 125 µM stocks of conjugation annealing handle and probe.
        ii. Add pre-annealed probe complex to nanoparticle pellet and add appropriate volumes of 1x and 10x wash buffer to achieve a final concentration of 1x wash buffer and a final volume 3.3x the initial nanoparticle volume. Incubate at 24 °C for at least 16 hours with shaking at 800 RPM on a ThermoMixer dry block. Suggested volumes:
            1. 40-nm particles: For a final volume of 200 µl, combine 176 µl probe complex, 17.6 µl 10x wash buffer, and 4 µl wash buffer.
            2. 200-nm particles: For a final volume of 20 µl, combine 17 µl probe complex, 1.7 µl 10x wash buffer, and 1.3 µl wash buffer.
    b. Post-anneal (optimized for 200-nm particles):
        i. Post-anneal the conjugation-annealing handle (see **Table 3** for details) to the particle by adding conjugation annealing handle to nanoparticle pellet and add appropriate volumes of 1x and 10x wash buffer to achieve a final concentration of 1x wash buffer and a final volume 3.3x the initial nanoparticle volume. Incubate at 24 °C for at least 16 hours with shaking at 800 RPM on a ThermoMixer dry block. Suggested volume:
            1. 200-nm particles: For a final volume of 20 µl, combine 8.5 µl conjugation-annealing handle, 0.85 µl 10x wash buffer, and 10.7 µl wash buffer.
        ii. Wash particles and redisperse in wash buffer equal to 28x the initial particle volume twice. Suggested volume:
            1. 200-nm particles: 180 µL.
        iii. Wash particles. Redisperse in a solution of 125 µM DNA aptamer probe at equal molar ratio to the conjugation-annealing handle. Bring to 3.3x the initial particle volume using water and incubate at 95 °C for 5 minutes. Centrifuge briefly to remove condensation from the tube lid. Incubate on benchtop for 10-15 minutes to allow the two complementary regions of DNA to anneal. Suggested volumes:

1. 200-nm particles: Combine 8.5 µl of 125 µM probe and 11.5 µl DNase/RNase free water.

10. Add 1 µL wash buffer/µL PEG solution for 40-nm particles and 7 µL wash buffer/µL PEG solution for 200-nm particles. Suggested volumes:
    a. 40-nm particles: 250 µL wash buffer.
    b. 200-nm particles: 180 µL wash buffer.
11. Wash and redisperse particles in 2 µL wash buffer/µL PEG solution PBS for 40-nm particles and 5 µL wash buffer/µL PEG solution for 200-nm particles. (Note: The pellet will be less defined, and some nanoparticles may remain in solution after this step.) Suggested volumes:
    a. 40-nm particles: 500 µL wash buffer.
    b. 200-nm particles: 125 µL wash buffer.
12. Repeat Step 11. Store nanoparticles in wash buffer at 4 °C.

***PEG Density Calculations***

The density of PEG-36 (MW of 1.6 kDa) required to achieve brush layer conformation was determined utilizing theoretical and experimental models[2–4]. The PEG conformation depends upon two parameters: 1) the Flory radius ($R_F$), which is the radius of the PEG coil and is dependent upon molecular weight and 2) the distance between PEG molecule graft sites (D). The relationship between $R_F$ and D dictates the PEG conformation: If $D > R_F$, the PEG layer will be a mushroom conformation; if $D < R_F$, it will be a brush layer; and if $D < 0.36\ R_F$ it will be a dense brush layer[2–4] . $R_F$ can be calculated using the following equations[2–4]:

$$R_F = \alpha N^{3/5}$$

where $\alpha$ = the length of one monomer and N = the number of monomers/polymer chain.

*Assumptions:*

- D = distance between PEG molecules
- $\alpha = 0.35$ nm
- N = 36
- Nanoparticle diameter = 200 nm
- Each PEG molecule occupies one circular area ($A_{PEG}$) on the particle surface

- PEG molecules are equally spaced across the surface
- A dense PEG layer requires $D < 0.36R_F$

*Calculations:*

- Flory Radius:

  $$\mathrm{R}_F = (0.35\, nm)\left(36^{\frac{3}{5}}\right) = 3.0\, nm$$

- Surface area of the nanoparticle:

  $$Surface\ area_{NP} = 4\pi r^2 = 4\pi(100nm)^2 = 125{,}663\ nm^2$$

- PEG distance to achieve a dense brush layer:

  $$D < (0.36)R_F, D < 1.08\, nm$$

- PEG distance as a function of PEG spacing:

  Distance between two PEG molecules:

  $$D = Radius\ of\ PEG1 + Radius\ of\ PEG2 = Diameter\ of\ A_{PEG}$$

  $$A_{PEG} = \pi(\frac{D}{2})^2$$

  $$D = 2\left(\frac{A_{PEG}}{\pi}\right)^{1/2}$$

  $$D < 1.08\, nm$$

  $$1.08\, nm > 2\left(\frac{A_{PEG}}{\pi}\right)^{\frac{1}{2}}$$

  $$A_{PEG} < 0.92\, nm^2$$

- Determination of minimum # PEG molecules/particle:

  $$A_{PEG} = \frac{Surface\ area_{NP}}{\#\ PEG\ molecules}$$

$$0.92\ nm^2 = \frac{125{,}663\ nm^2}{\#\ PEG\ molecules}$$

$$\#\ PEG\ molecule > 137{,}000, or\ at\ least\ 10^5 PEG\ molecules/particle$$

**Supplemental References**


1. Pipkorn, R., Boenke, C., Gehrke, M. & Hoffmann, R. High-throughput peptide synthesis and peptide purification strategy at the low micromol-scale using the 96-well format. *J. Pept. Res.* **59**, 105–114 (2002).

2. Yang, Q. *et al.* Evading immune cell uptake and clearance requires PEG grafting at densities substantially exceeding the minimum for brush conformation. *Mol. Pharm.* **11**, 1250–1258 (2014).

3. Damodaran, V. B., Fee, C. J., Ruckh, T. & Popat, K. C. Conformational Studies of Covalently Grafted Poly ( ethylene glycol ) on Modified Solid Matrices Using X-ray Photoelectron Spectroscopy. **26**, 7299–7306 (2010).

4. Perry, J. L. *et al.* PEGylated PRINT Nanoparticles: The Impact of PEG Density on Protein Binding, Macrophage Association, Biodistribution, and Pharmacokinetics. (2012).

**Supplemental Figures and Figure Legends**

**Figure S1**

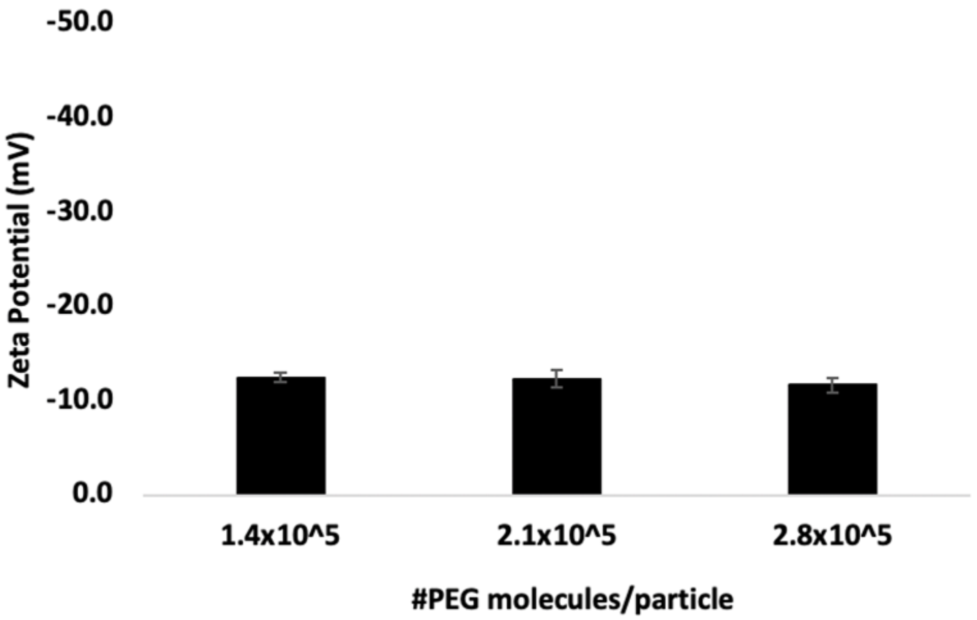


**Supplementary Figure S1. PEG density for 40-nm nanoparticles.** Zeta potential measurements for 40-nm carboxylated FluoSpheres™ activated with NHS/EDC and reacted with increasing concentrations of mPEG-amine. Data is depicted as means ± standard deviation of the six measurements taken (three per replicate), n=2.

**Figure S2**

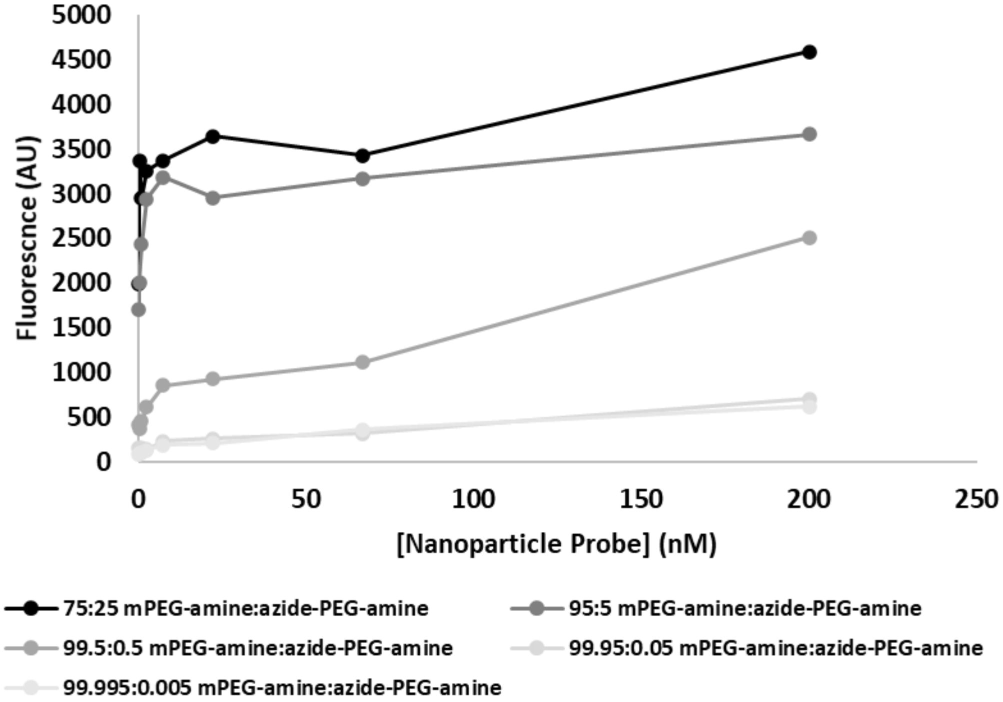


**Supplementary Figure S2. On-target binding of nanoparticle B1 probes fabricated with various ratios of mPEG-amine to azide-PEG-amine.** Binding of fluorescent nanoparticle B1 probes with varying ratios of mPEG-amine to azide-PEG-amine to his-tagged Her2 protein. Data are from a single experiment.

**Figure S3**

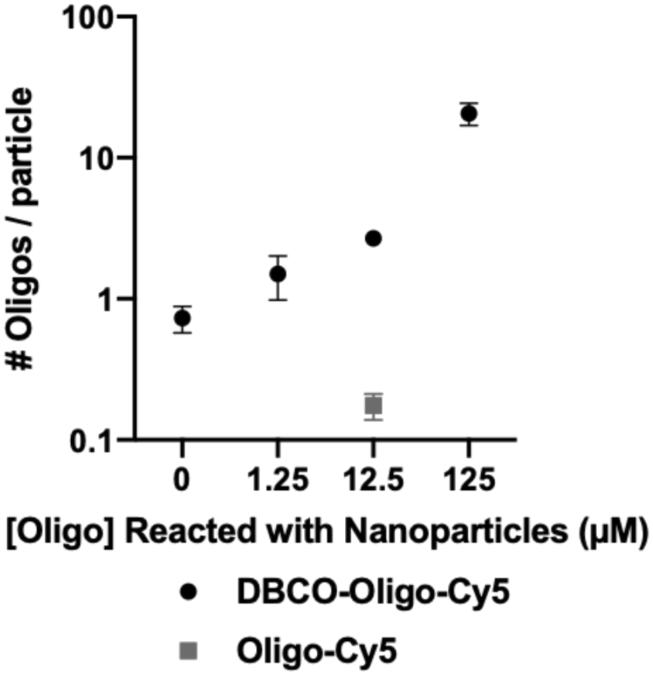


**Supplementary Figure S3. Conjugation of the conjugation-annealing handle to fluorescent nanoparticles.** Ratio of detected oligos per nanoparticles for PEGylated nanoparticles reacted with 0, 1.25, 12.5, and 125 µM of a fluorescently-labeled conjugation-annealing handle ("DBCO-oligo-Cy5"). PEGylated nanoparticles were reacted with 12.5 µM of the fluorescently-labeled conjugation-annealing handle without the DBCO group ("oligo-Cy5") as a non-reactive control. Data graphed is depicted as means ± standard deviation, n=3-4.

**Figure S4**

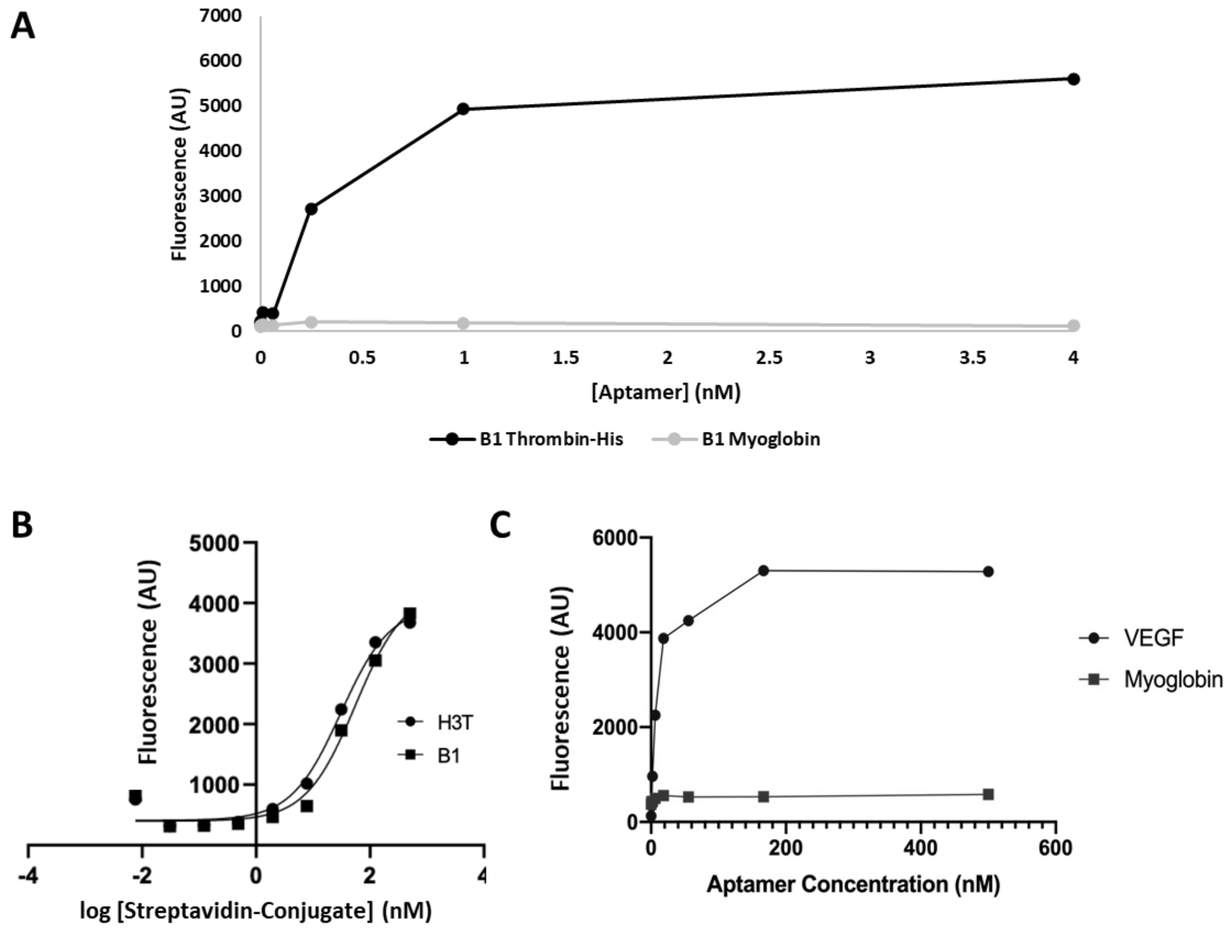


**Supplementary Figure S4. Binding of aptamers detected with streptavidin-HRP or streptavidin-conjugates.** (A) Binding of a biotinylated B1 aptamer to his-tagged thrombin detected by streptavidin-HRP. (B) Binding of B1 and H3T streptavidin conjugates to HHH peptide, with calculated $EC_{50}$s of 55 and 30 nM, respectively. (C) Binding of a biotinylated VEGF aptamer to VEGF protein detected by streptavidin-HRP.

**Figure S5**

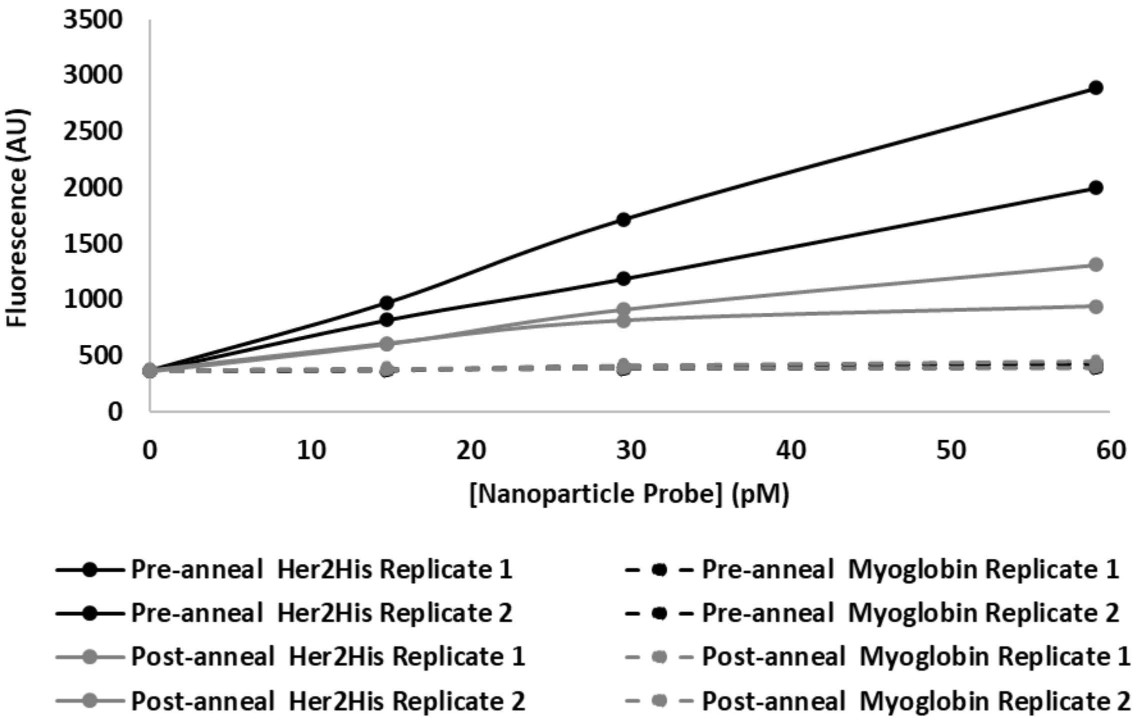


**Supplementary Figure S5. Affinities of probes prepared by pre- and post-annealing of DNA aptamers to nanoparticles.** Binding of fluorescent nanoparticle B1 probes fabricated with pre- and post-annealing protocols to his-tagged Her2 (on-target) and myoglobin (off-target). Data are representative of a single experiment performed in duplicate.

**Figure S6**

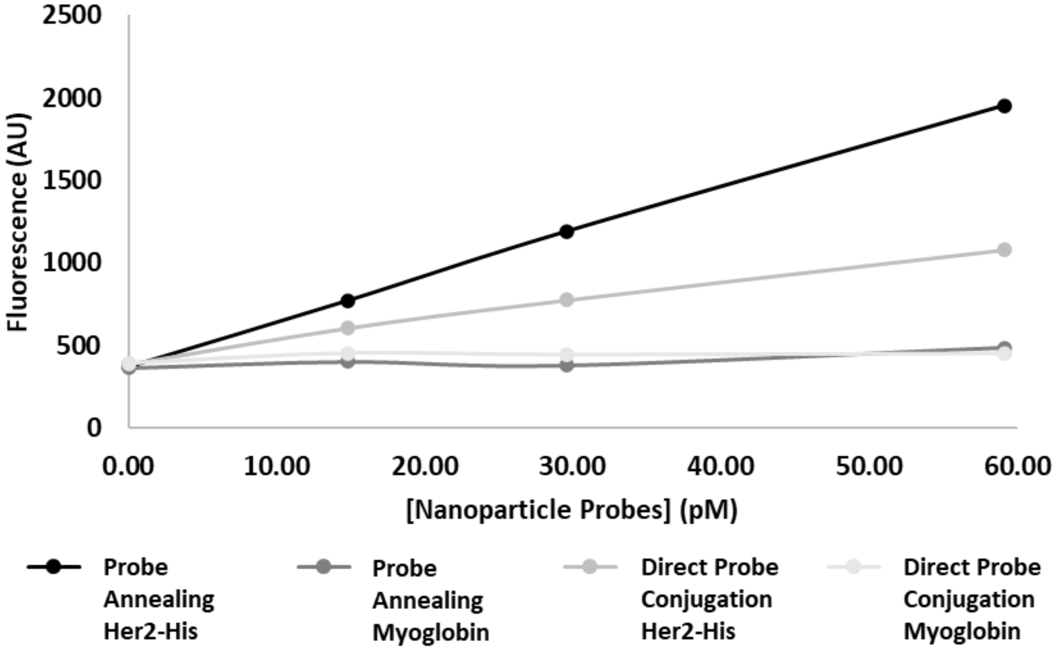


**Supplementary Figure S6. Affinity of probe prepared by direct conjugation of DNA aptamer to nanoparticles.** Binding of fluorescent nanoparticle B1 probes fabricated with direct conjugation or with pre-annealing protocol to his-tagged Her2 (on target) and myoglobin (off-target). Data are from a single experiment.

**Figure S7**

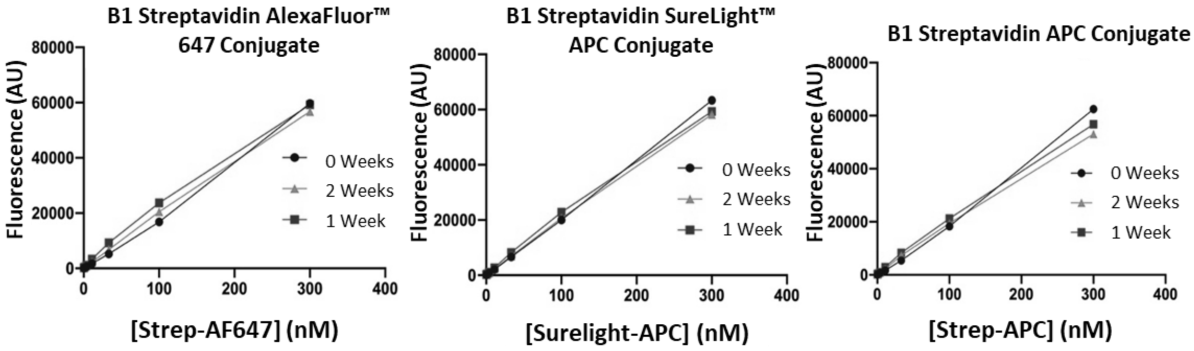


**Supplementary Figure S7. Fluorescent intensities of labels are stable over 2 weeks.**

Fluorescent intensity readings of commercially available labels conjugated to the B1 aptamer measured over the course of two weeks.

**Figure S8**

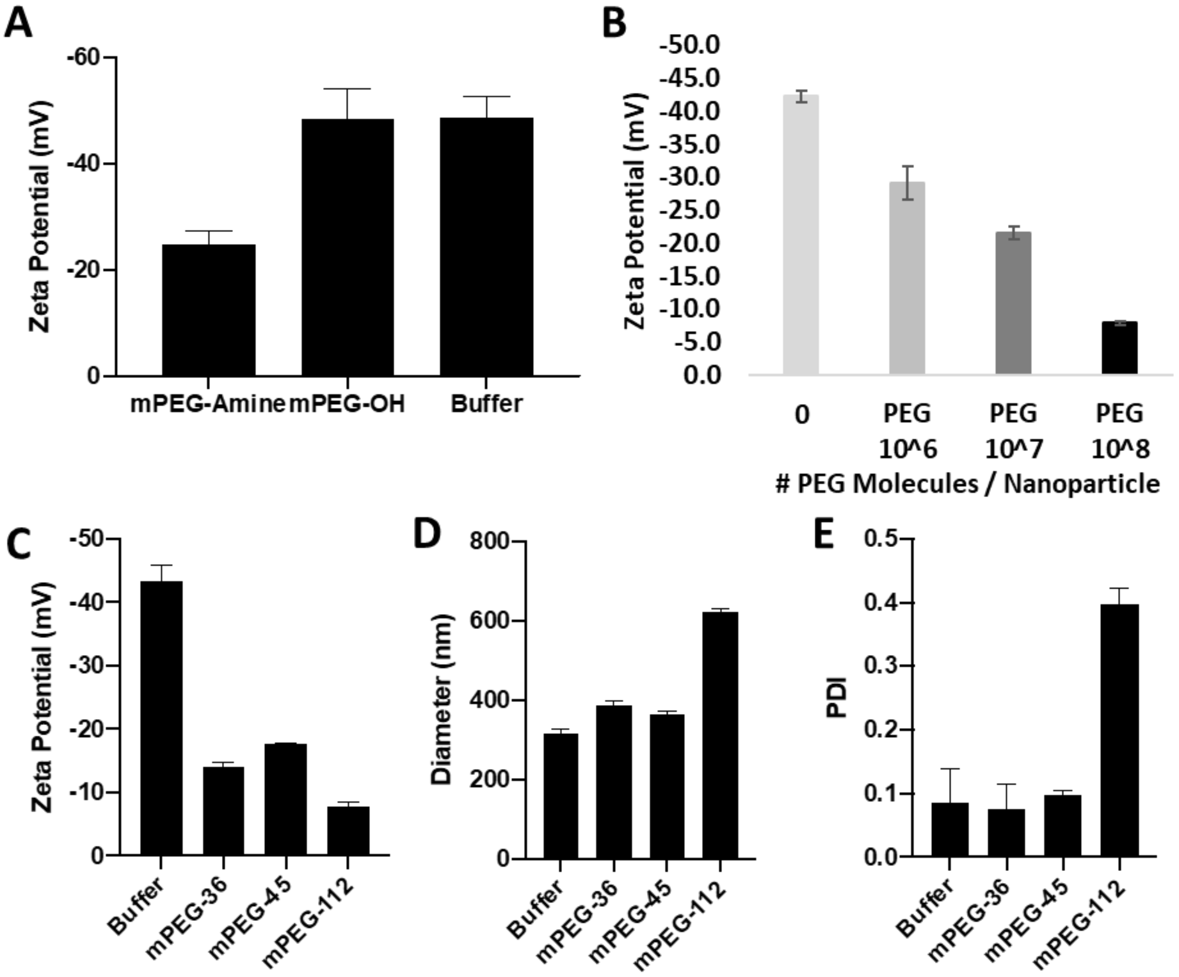


**Supplementary Figure S8. Replicates for Figure 2 PEGylation of carboxylate-modified FluoSpheres™.** (A) Zeta potential measurements for carboxylated FluoSpheres activated with NHS/EDC and reacted with mPEG-amine, mPEG-OH, or buffer only. Data are means ± standard deviation of measurement replicates, n=2. (B) Zeta potential measurements for carboxylated FluoSpheres activated with NHS/EDC and reacted with increasing concentrations of mPEG-amine. Data are means ± standard deviation, n=3. (C-E) FluoSpheres were conjugated with PEGs of indicated molecular weights, and C) zeta potentials, (D) hydrodynamic diameters, and (E) PDI were determined. Data are means of measurement replicates, n=1-2 experimental replicates.

### Figure S9

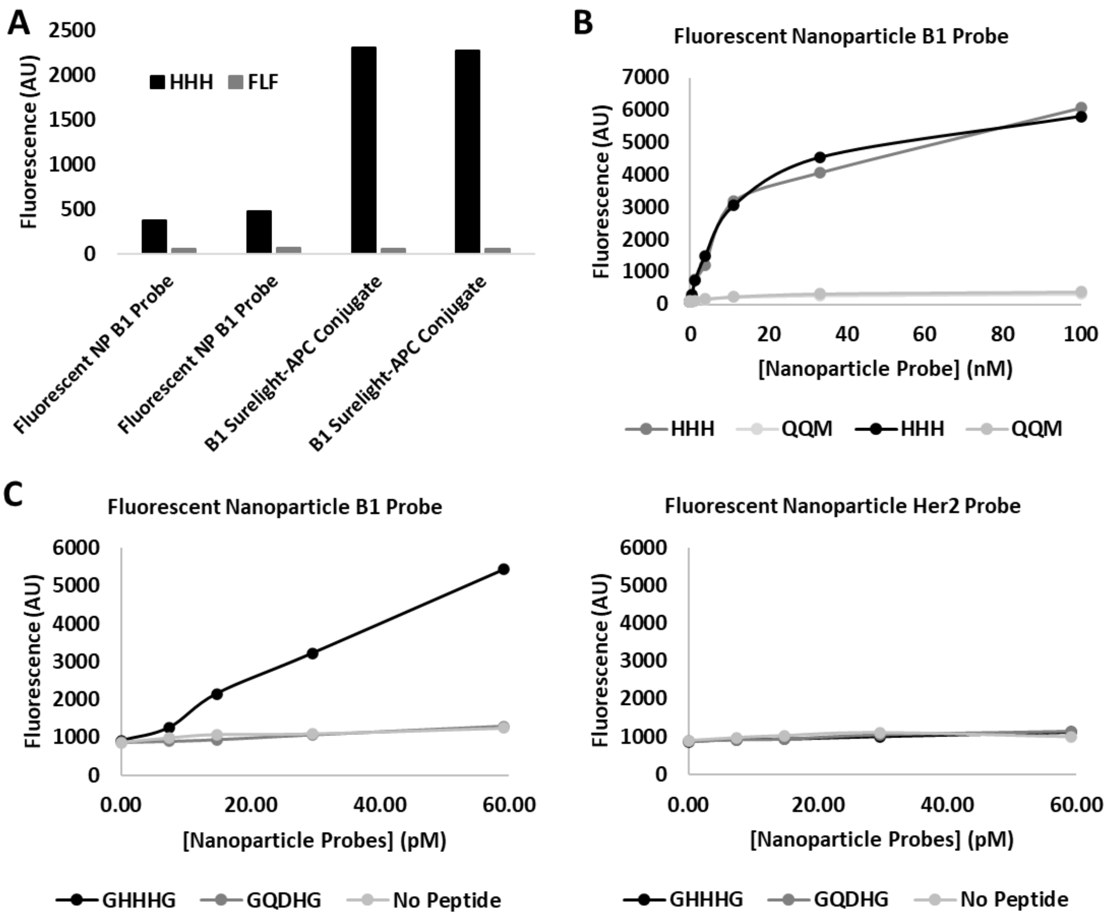


**Supplementary Figure S9. Replicates for Figure 4 Fluorescent nanoparticle B1 probe binds specifically to HHH peptide.** (A) Fluorescence signals from fluorescent nanoparticle B1 probes and B1 Surelight-APC conjugates against HHH (on-target) or FLF (off-target) peptides. Data shows duplicates from a single experiment. (B) Fluorescence of B1 probe as a function of concentration against HHH peptide (on-target) and QQM peptide (off-target). Data shows duplicates from a single experiment. (C) Fluorescence of B1 and Her2 probe fluorescent nanoparticle probes against HHH and QDH peptides. Data is from a single experiment.

**Figure S10**

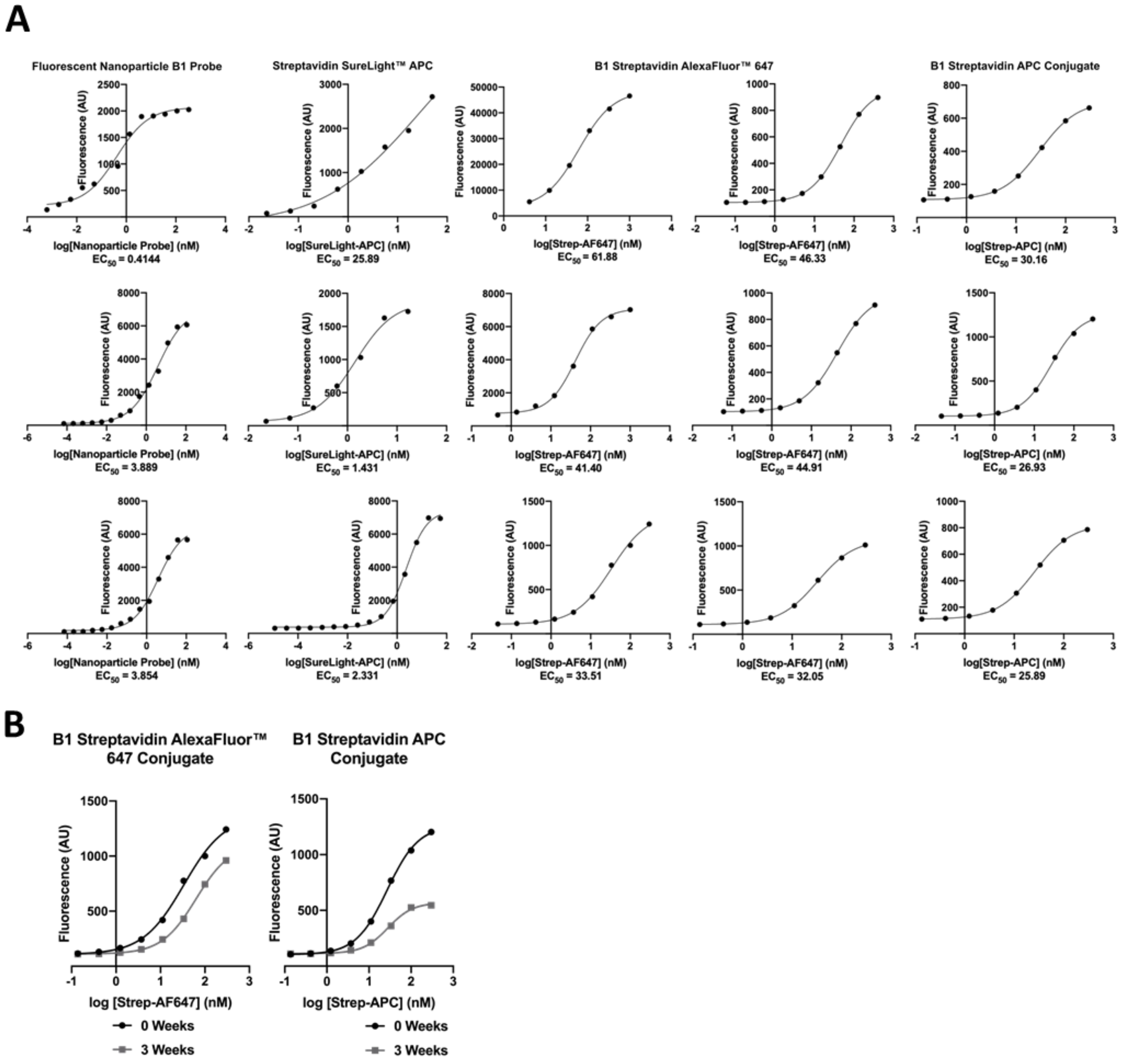


**Supplementary Figure S10. Replicates for Figure 6 Comparison of Fluorescent nanoparticle probes with commercially available labels.** (A) Binding curves and $EC_{50}$ values for fluorescent nanoparticle B1 probe, B1 Streptavidin SureLight™ APC, B1 Streptavidin AlexaFluor™ 647 Conjugate, and B1 Streptavidin APC Conjugate against HHH targets. These binding curves were used to create the box and whiskers plot shown in Figure 6A, and one representative graph per probe from this panel is shown in Figure 6A. (B) Fluorescence intensities from binding curves of B1 Streptavidin AlexaFluor™ 647 Conjugate and B1 Streptavidin APC Conjugate against HHH targets at noted timepoints post fabrication.

**Figure S11**

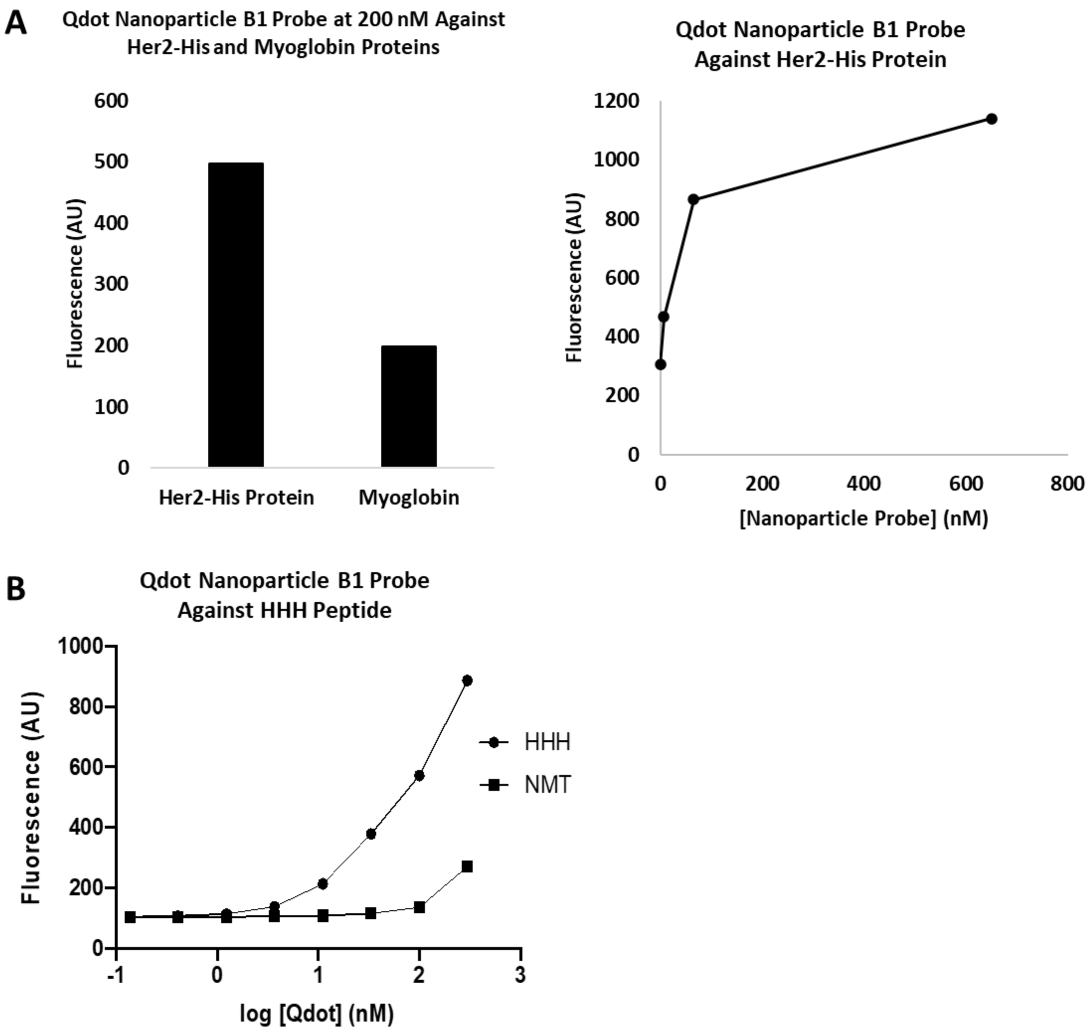


**Supplementary Figure S11. Replicates for Figure 7 Fluorescent nanoparticle probes with a Quantum Dot core.** Qdot™ 655 ITK™ Carboxyl Quantum Dots from Thermo Fisher Scientific were used as an alternative to FluoSpheres™ as the nanoparticle core. (A) Binding data and curves of the Qdot nanoparticle B1 probe to Her2-his (on-target) and myoglobin (off-target) proteins. Data shown is means of two duplicates in one experiment for both graphs. (B) Binding curves of the Qdot fluorescent nanoparticle B1 probe to HHH peptide. Data shown is means of two duplicates in one experiment.

**Figure S12**

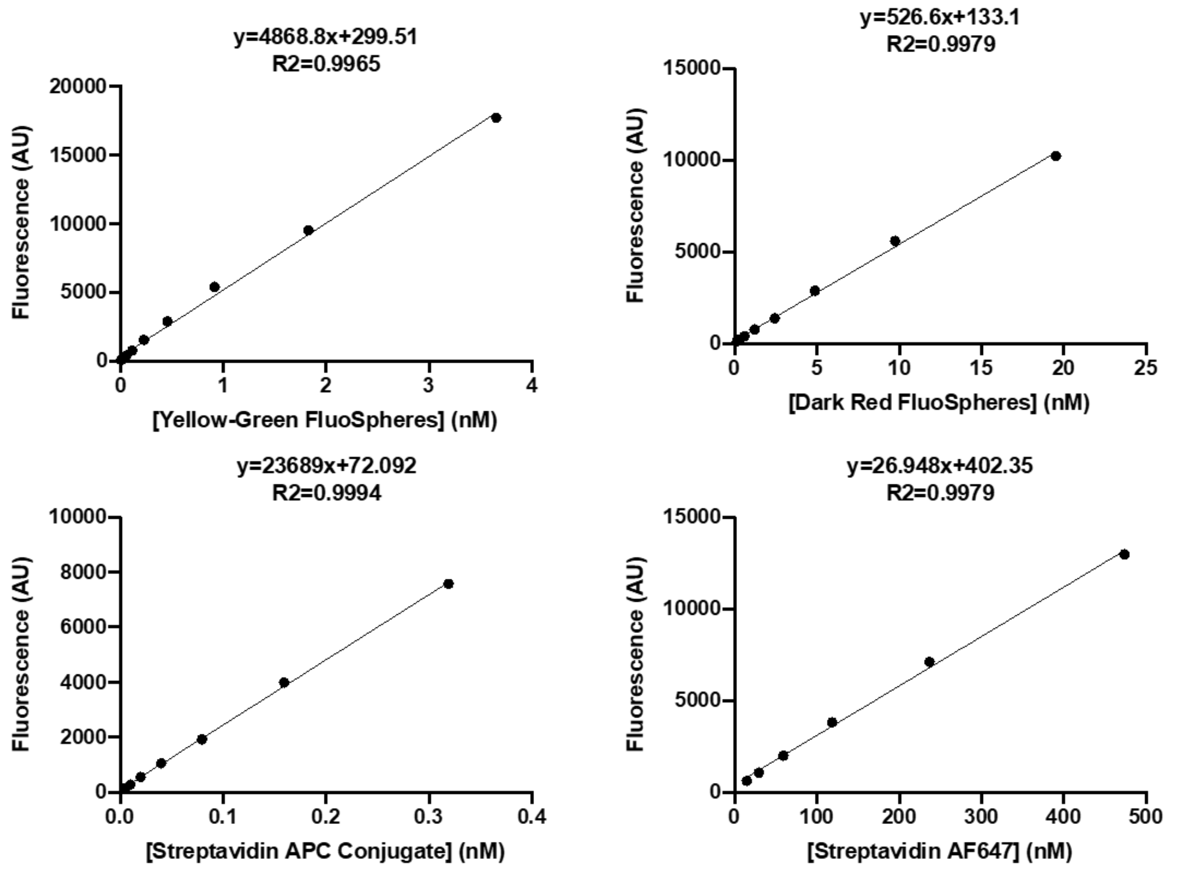


| Label | Upper Label Concentration | Lower Label Concentration |
|---|---|---|
| Yellow-Green FluoSpheres™ | 3.65 nM | 0.007 nM |
| Dark Red FluoSpheres™ | 19.5 nM | 0.15 nM |
| Streptavidin APC Conjugate | 319 nM | 4.98 nM |
| Streptavidin AlexaFluor™ 647 | 474 nM | 14.8 nM |

| Wavelength | Upper Limit Fluorescence | Lower Limit Fluorescence |
|---|---|---|
| 485/535 | 17,725 AU | 57 AU |
| 635/680 | 10,000 - 13,000 AU | 100 - 600 AU |

**Supplementary Figure S12. Limit of detection for labels on plate reader.** The limit of detection for each label was determined for the fluorescent plate reader used in binding assays. The linear fit is shown for each label. The charts summarize the upper and lower concentrations limits for each label as well as the upper and lower fluorescent limits for each wavelength measured.
